# The ACDC domains of AP2-O3 and FD3 contribute to protein interactions and ookinete development

**DOI:** 10.64898/2026.11.19.767835

**Authors:** Jingjing Tang, Bing Liu, Aniko Verbrugge, Yangxin Xie, Isadora Oliveira Prata, Manuel Llinás, Zhenkui Li

**Affiliations:** Institute of Pathogenic Biology and Key Laboratory of Special Pathogen Prevention and Control of Hunan Province, School of Basic Medical Sciences, Hengyang Medical School, University of South China, Hengyang, 421001, China; Zhejiang Provincial Key Laboratory of Precision Diagnosis and Therapy for Major Gynecological Diseases, Women’s Hospital, Zhejiang University School of Medicine, Hangzhou, 310058, China; Department of Biochemistry and Molecular Biology, Pennsylvania State University, University Park, PA 16802, USA; Huck Center for Malaria Research, Pennsylvania State University, University Park, PA 16802, USA; Department of Chemistry, Pennsylvania State University, University Park, PA 16802, USA; Department of Human Parasitology, School of Basic Medical Sciences, Hengyang Medical School, University of South China, Hengyang, 421001, China

**Keywords:** *Plasmodium*, ACDC domain, malaria transmission, ApiAP2 transcription factors, sexual development

## Abstract

The Apicomplexan AP2 (ApiAP2) family of transcription factors are key regulators of *Plasmodium* spp. parasite development. In addition to containing AP2 DNA-binding domains, nine ApiAP2 proteins also encode an apicomplexan-specific ACDC domain whose function remains largely unknown. Here, we dissect the biological role of the ACDC domain from two female-specific regulators, AP2-O3 and FD3, using the rodent malaria model *Plasmodium yoelii* and the human malaria parasite *P. falciparum*. In *Plasmodium yoelii*, deletion of the ACDC domain preserved AP2-O3 and FD3 protein presence and nuclear localization without significantly affecting asexual growth or gametocyte production. However, deletion of these domains impaired the gametocyte-to-ookinete transition, reducing ookinete conversion efficiency. Proteomic and biochemical studies indicate that the AP2-O3 ACDC domain interacts with FD3, while the FD3 ACDC domain preferentially associates with full-length FD3. Transcriptomic data showed that over 80% of transcripts that increased after FD3 ACDC deletion were also higher following AP2-O3 loss, with female gametocyte marker expression largely unchanged. In *Plasmodium falciparum*, removing the FD3 ACDC domain did not affect female lineage-specific expression of FD3, nuclear localization of FD3, or the asexual growth and gametocyte maturation of parasites. Leveraging the high sequence conservation of ACDC domains across the *Plasmodium* genus, we conducted structure-based virtual screening of 283 compounds targeting the predicted PfAP2-O3 and PfFD3 ACDC interaction interface. Seven compounds ranked among the top candidate inhibitors for both domains, and the highest-priority ligand for each domain was further evaluated by molecular dynamics simulations. Collectively, our findings support a role for ACDC domains in ApiAP2 protein interactions and identify candidate compounds for future experimental evaluation as potential transmission-blocking leads.

## Introduction

Malaria continues to be a significant global health challenge, causing hundreds of thousands of deaths each year despite ongoing control measures [1]. The expansion of antimalarial drug resistance, combined with the lack of effective transmission-blocking strategies, continues to hinder efforts to eradicate malaria [2]. Parasite transmission from the mammalian host to the mosquito host relies on a precisely coordinated developmental transition from gametocytes to motile ookinetes [3, 4], a process accompanied by extensive transcriptional remodeling. Nevertheless, the molecular mechanisms that govern stage-specific transcriptional programming during this critical transition remain incompletely characterized [5].

Unlike most eukaryotes, which utilize diverse transcription factor families, *Plasmodium* parasites depend heavily on the ApiAP2 (Apicomplexan AP2) family for developmental regulation throughout their life cycle [6]. ApiAP2 proteins possess one or more AP2 DNA-binding domains, are evolutionarily derived from the plant APETALA2/ERF domain, and are thought to have been acquired early in apicomplexan evolution [7]. *Plasmodium* genomes encode approximately 30 ApiAP2 members [8], many of which act as stage-specific molecular switches. By binding cognate DNA motifs in promoter regions [9], ApiAP2 proteins can activate or repress transcriptional programs that govern asexual replication in hepatocytes and erythrocytes, as well as sexual development in the mosquito [8, 10–12]. Understanding how this restricted set of ApiAP2 factors produces such a wide range of regulatory outputs required for parasite development, beyond just recognizing DNA sequences, remains a crucial unanswered question.

Nine ApiAP2 proteins harbor an additional highly conserved region designated as the ACDC (AP2-Coincident Domain, mostly at the C-terminus), an apicomplexan-specific module absent from mammalian hosts and evolutionary progenitors such as the green-brown algae *Chromera spp* [13]. Recent structural work has demonstrated that the ACDC domain adopts a distinctive four-helix bundle fold with no detectable homology to known eukaryotic regulatory domains [14]. ACDC-containing ApiAP2 factors have been linked to a broad range of developmental processes, including liver-stage progression [15], erythrocyte invasion [16], sexual commitment [17, 18], and mosquito-stage development [10, 11]. Despite the importance of ApiAP2 proteins to parasite development, the ACDC domain has no established molecular function. Whether it acts directly in transcriptional regulation or recruits protein partners to confer regulatory specificity remains unknown.

The AP2-O3 ApiAP2 protein contains an ACDC domain and is a female-specific transcriptional repressor that preserves the female transcriptome by repressing male-associated genes [19]. Disruption of *ap2-o3* (PY17X_1017000) in *Plasmodium yoelii* causes ectopic expression of male genes in female gametocytes and impairs female gametogenesis and proper ookinete formation [19]. FD3 (PBANKA_1418100), another ACDC-containing ApiAP2 protein, has been identified as a transcriptional activator that promotes the expression of female-specific transcripts in *Plasmodium berghei* [20]. Additionally, the AP2-O3 ortholog (*Pb*AP2-FG2, PBANKA_1015500) has been reported to function as part of a transcriptional repressor complex with FD3 (*Pb*AP2R-2) in *P. berghei* female gametocytes [21]. While these studies indicate that protein-protein interactions between AP2-O3 and FD3 mediate ApiAP2 regulation, it remains unclear how this occurs and whether the ACDC domain facilitates these associations.

In this study, we investigated the role of the ACDC domains of AP2-O3 and FD3 in protein-protein interactions and transmission-stage development. Our results show that the AP2-O3 and FD3 proteins interact in *P. yoelii* gametocytes and that the isolated ACDC domains recover overlapping but distinct sets of proteins from gametocyte lysates. Deleting the ACDC partially hinders ookinete conversion, although it does not affect the core transcriptional repertoire of gametocytes. We also predicted ACDC structures to conduct structure-based virtual screening of a compound library, prioritizing potential ligands that may disrupt protein interactions mediated by ACDC domains. Overall, our results demonstrate the importance of ACDC-mediated complex assembly in ApiAP2-dependent regulation of malaria parasite development and suggest candidate ligands for further research.

## Results

### Deletion of the AP2-O3 ACDC impairs ookinete development

To investigate the functional contribution of the ACDC domain from the female-specific transcriptional regulator AP2-O3, we generated endogenous *ap2-o3* modifications in *Plasmodium yoelii*. PyAP2-O3 contains one N-terminal AP2 DNA-binding domain and one C-terminal ACDC domain. We first generated an N-terminally 6HA-tagged parental line via CRISPR/Cas9-mediated homologous recombination, *6HA::ap2-o3,* and verified correct integration by genomic junction PCR (Fig. S1A, B). From this parental line, we introduced a scarless in-frame deletion of the ACDC-coding region, yielding the *6HA::ap2-o3;ap2-o3Δacdc* strain (Fig. S1C-E). Separately, an untagged *ap2-o3Δacdc* line was generated from wild-type *P. yoelii* 17XNL. We measured the presence of the *Py*AP2-O3 protein and its subcellular localization in gametocytes by western blot and immunofluorescence microscopy. Anti-HA immunoblotting detected full-length AP2-O3 in the parental strain and a smaller band in *6HA::ap2-o3;ap2-o3Δacdc*, corresponding to the expected size of the ACDC-truncated protein (Fig. 1A). Immunofluorescence microscopy showed that ACDC deletion did not alter the nuclear localization pattern of AP2-O3 in female gametocytes (Fig. 1B). We then evaluated the developmental consequences of ACDC deletion across multiple parasite stages. The previously characterized *Δap2-o3* line was included as a loss-of-function comparator [19]. Asexual blood-stage growth over a 10-day time course revealed comparable parasite growth kinetics and multiplication rates among WT, *Δap2-o3*, and *ap2-o3Δacdc* parasites (Fig. 1C). Gametocytemia and male-to-female sex ratios were likewise similar across the three lines (Fig. 1D). In vitro ookinete conversion assays showed that WT parasites efficiently formed mature banana-shaped ookinetes. In contrast, *Δap2-o3* parasites failed to produce mature ookinetes, as previously reported [10, 11, 19]. The *ap2-o3Δacdc* mutant exhibited a significant but partial reduction in mature ookinete conversion efficiency relative to WT controls (Fig. 1E).

**Figure 1.**
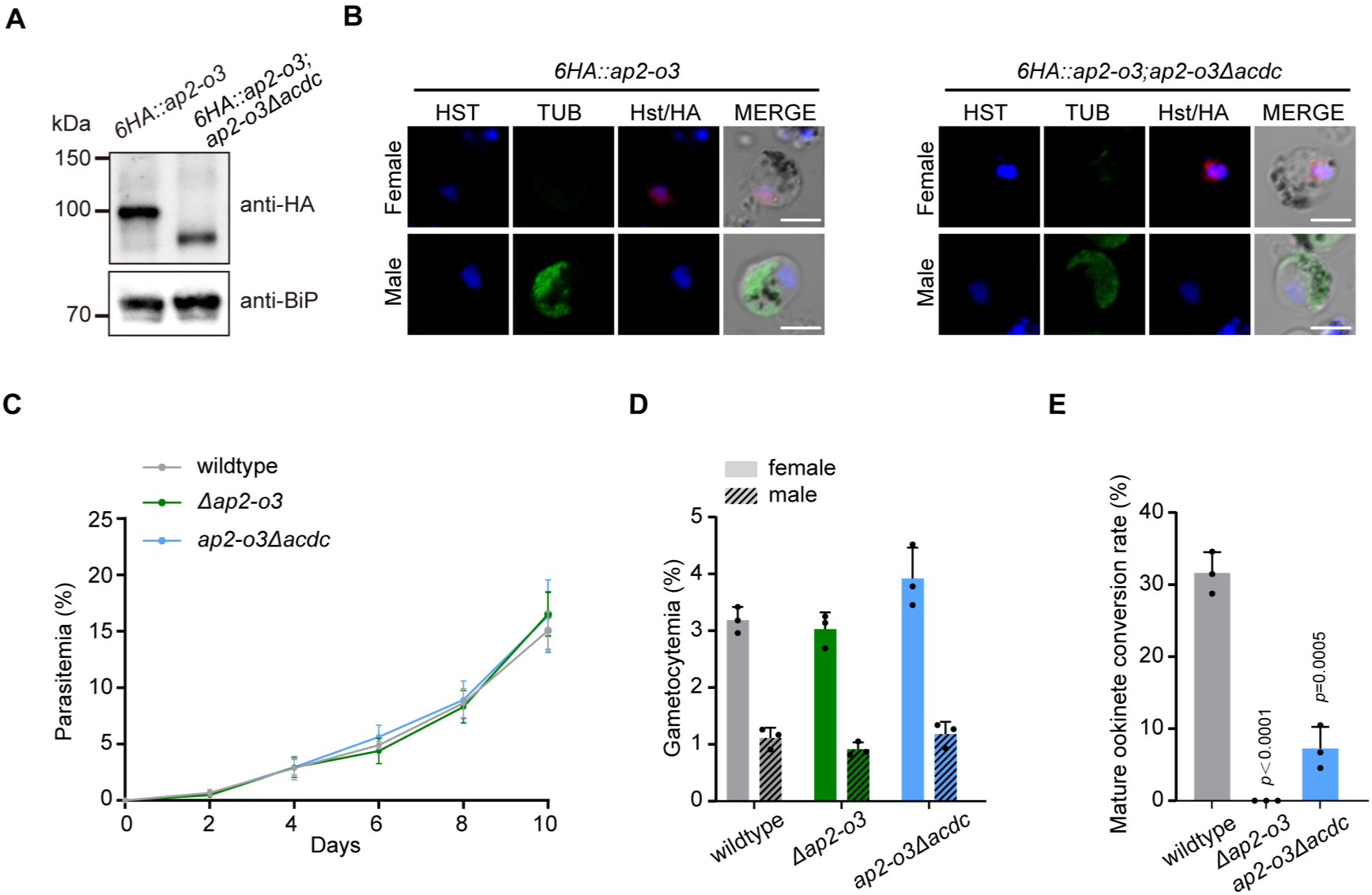
The AP2-O3 ACDC domain is not necessary for protein expression or nuclear localization but is required for effective ookinete development. (A) Western blot showing AP2-O3 levels in gametocyte lysates from various *P. yoelii* lines, detected with anti-HA antibody. BiP was used as a loading control. Results represent three independent biological replicates. (B) Immunofluorescence microscopy of AP2-O3 localization in female and male gametocytes from the *6HA::ap2-o3* parental and *6HA::ap2-o3;ap2-o3Δacdc*. AP2-O3 was visualized with anti-HA antibody; tubulin and Hoechst 33342 stain the male gametocyte and nuclei, respectively. Scale bar=5 μm. (C) Growth of WT, *Δap2-o3*, and *ap2-o3Δacdc* parasites during 10 days of asexual blood stages, assessed via Giemsa-stained thin blood smears. Data are mean ± SD from 3–5 mice per group. (D) Quantification of male and female gametocytemia in WT and mutant lines, expressed as mean ± SD from three mice per group. (E) In vitro development of ookinetes after 20 hours of culture, with mature ookinetes counted per 100 female gametocytes. Results are mean ± SD from three biological replicates. Statistical significance was determined by two-tailed Student’s t-test; P values are indicated in the graph.

### The AP2-O3 ACDC domain interacts with FD3

We next asked whether the ACDC domain of *Py*AP2-O3 mediates this interaction. A GST-tagged *Py*AP2-O3 ACDC domain was expressed and purified. As a matched negative-control bait, we prepared in parallel an identically purified GST fusion of the compact LOTUS domain (residues 144–263) of the gamete egress protein GEP (PY17X_1116400), an unrelated parasite protein that is not expected to participate in ACDC-mediated interactions. Both baits were incubated with *P. yoelii* gametocyte lysates in GST pull-down assays, followed by LC-MS/MS analysis (Fig. 2A, B). Across three independent experiments, and applying stringent enrichment thresholds for the ACDC bait relative to the GST–LOTUS control, we identified 80 proteins significantly enriched by the *Py*AP2-O3 ACDC domain pull-down, including FD3 (Fig. 2C, Table S1). Peptide-level analysis showed that all 10 unique FD3-derived peptides identified in the AP2-O3 ACDC pull-down mapped to the ACDC region of FD3 (Table S1). In addition to FD3, the enriched set also contained a subset of nuclear proteins including core histones (H2A, H2B, and H4), a high-mobility-group protein, and spliceosome components (Table S1). Other proteins detected include ribosomal, translation, and cytoskeletal proteins that may reflect indirect or nonspecific associations that require targeted validation. GST pull-down coupled with immunoblotting was used to validate the interaction between FD3 and the *Py*AP2-O3 ACDC domain (Fig. 2D). To test whether the AP2-O3 and FD3 proteins interact in *P. yoelii* gametocytes, we generated a dual-tagged parasite line (*6HA::ap2-o3; 4Myc::fd3*) (Fig. 2E). Co-immunoprecipitation with anti-Myc nanobody agarose beads enriched *Py*AP2-O3, as detected by anti-HA immunoblotting (Fig. 2F), confirming the AP2-O3–FD3 interaction in *P. yoelii* gametocytes. Together, these results show that AP2-O3 and FD3 are associated in *P. yoelii* gametocytes and that the isolated *Py*AP2-O3 ACDC domain enriches FD3 from gametocyte lysates.

**Figure 2.**
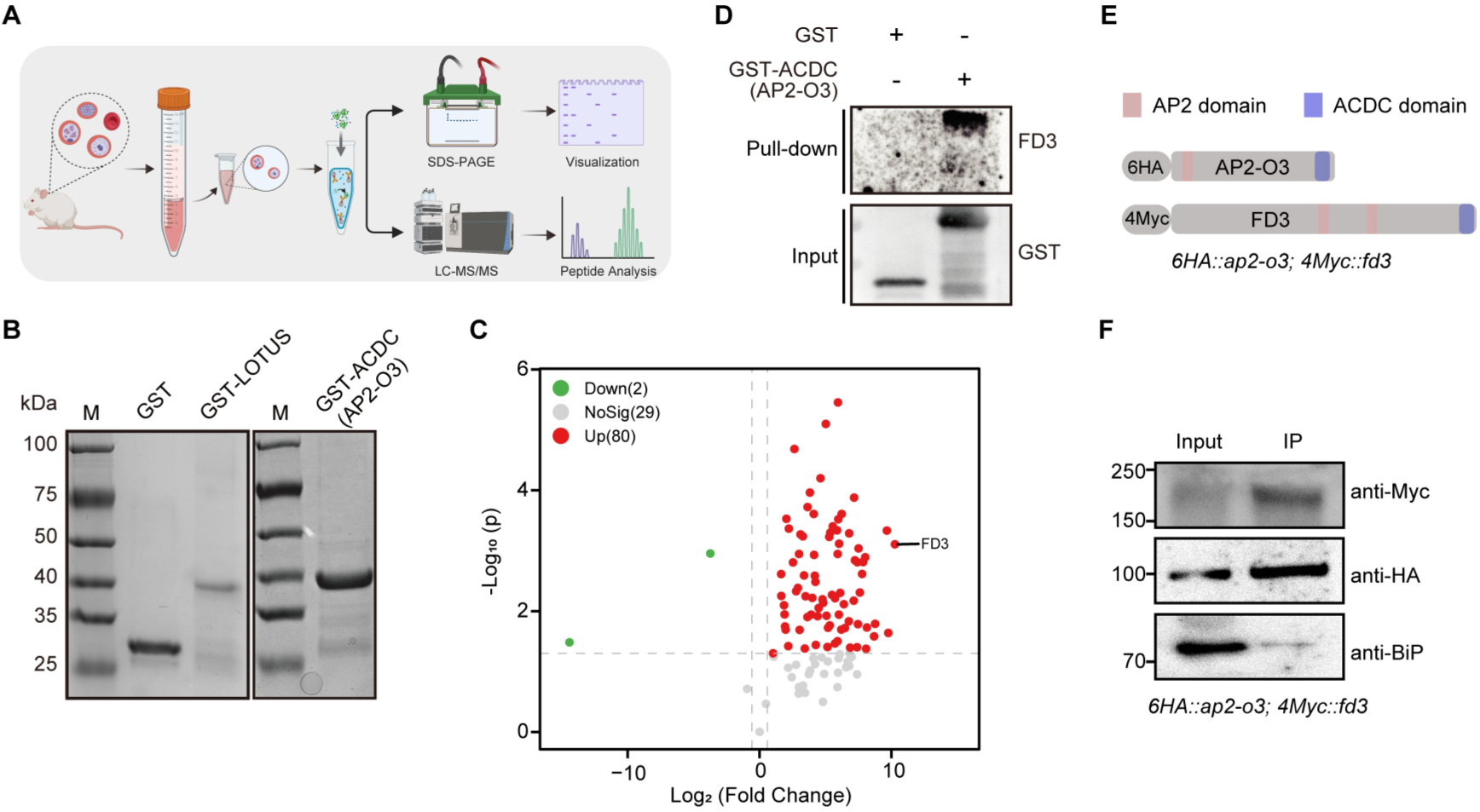
The AP2-O3 ACDC domain interacts with FD3 in *P. yoelii* gametocytes. **(A)** Workflow schematic for GST pull-down combined with LC–MS/MS to identify proteins associated with recombinant ACDC domains. **(B)** SDS-PAGE analysis of purified GST–AP2-O3– ACDC fusion protein. Empty GST vector was used as a control for the purification process, and GST-LOTUS served as the control for the subsequent GST pull-down coupled with mass spectrometry assay. **(C)** Volcano plot showing proteins enriched in GST–AP2-O3–ACDC pull-downs compared to GST–LOTUS negative-control pull-downs. Proteins with log₂ fold change ≥1 and P < 0.05 are highlighted; FD3 is among the enriched proteins. **(D)** Validation of the AP2-O3 ACDC–FD3 interaction through GST pull-down and immunoblotting. **(E)**Schematic of the dual-tagged parasite line (*6HA::ap2-o3; 4Myc::fd3*) used for analyzing protein interactions. The AP2 and ACDC domains are marked. **(F)** Co-immunoprecipitation of endogenous AP2-O3 and FD3 from gametocyte lysates of the *6HA::ap2-o3; 4Myc::fd3* line using anti-Myc nanobody agarose beads. Immunoprecipitates and input lysates were examined by immunoblot with anti-HA and anti-Myc antibodies; BiP was used as a loading control.

### The FD3 ACDC domain supports ookinete development and mediates FD3 oligomerization

*Py*FD3 is a 1,773-amino-acid protein containing a single C-terminal ACDC domain (residues 1683–1770) [22] and two cryptic AP2 domains [13] (Fig. S2). To assess the functional contribution of the FD3 ACDC domain, we generated *P. yoelii* mutants lacking this region. We first established an N-terminally 4Myc-tagged FD3 parental line and introduced an in-frame deletion of the ACDC-coding sequence using CRISPR/Cas9, yielding the *4Myc::fd3;fd3Δacdc* strain (Fig. S1F-H). We also generated an *fd3* knockout strain (*Δfd3*) as a loss-of-function control (Fig. S1I, J). Immunoblotting detected full-length FD3 in the parental line and a smaller band in *4Myc::fd3;fd3Δacdc*, corresponding to the expected size of ACDC-truncated FD3 (Fig. 3A). Both full-length and truncated FD3 localized to the nuclei of female gametocytes (Fig. 3B). We next compared asexual growth, gametocyte production, and ookinete conversion among WT, *Δfd3*, and *fd3Δacdc* parasites. All three lines exhibited comparable asexual replication and sexual commitment, with no significant differences in male or female gametocyte numbers (Fig. 3C, D). During *in vitro* ookinete development, *Δfd3* parasites completely failed to form mature ookinetes. At the same time, *fd3Δacdc* mutants showed a strong reduction in mature ookinete conversion rate relative to WT (Fig. 3E), similar to that observed in *ap2-o3Δacdc* mutants (Fig. 1E), supporting a possible heterodimer function.

**Figure 3.**
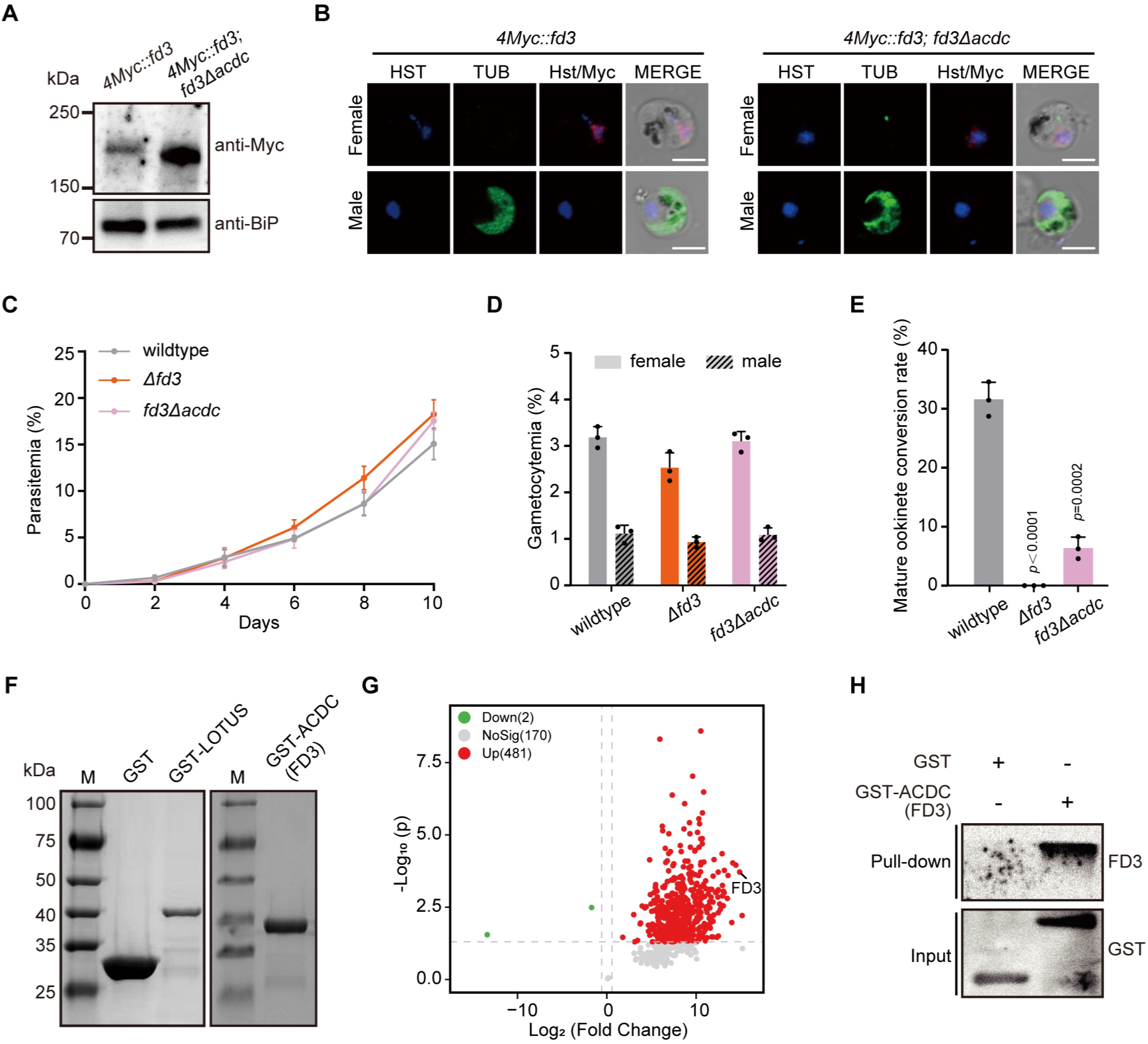
The FD3 ACDC domain promotes efficient ookinete development and mediates FD3-associated protein interactions. **(A)** Western blot analysis shows FD3 expression in gametocyte lysates from *4Myc::fd3* and *4Myc::fd3;fd3Δacdc* parasites, using anti-Myc antibody; BiP is a loading control. Results are representative of three independent experiments. **(B)** Immunofluorescence reveals FD3 localization in female and male gametocytes of both *4Myc::fd3* and *4Myc::fd3;fd3Δacdc* lines. FD3 was visualized with anti-Myc antibody; tubulin and Hoechst 33342 stain the male gametocyte and nuclei, respectively. Scale bar=5 μm. **(C)** Growth of WT, *Δfd3*, and *fd3Δacdc* parasites during the asexual blood stage was monitored over 10 days via Giemsa-stained blood smears; data represent means ± SD from 3–5 mice per group. **(D)** Gametocytemia levels in males and females were quantified in WT and mutant lines; data are means ± SD from three mice per group. **(E)** In vitro development of ookinetes after 20 hours of culture, with mature ookinetes counted per 100 female gametocytes; data show means ± SD from three biological replicates. Significance determined by two-tailed Student’s t-test. **(F)** SDS-PAGE analysis of purified GST– FD3–ACDC fusion protein. Empty GST vector was used as a control for the purification process, and GST-LOTUS served as the control for the subsequent GST pull-down coupled with mass spectrometry assay. **(G)** Volcano plot displays proteins enriched in GST–FD3–ACDC pull-downs relative to GST–LOTUS negative-control pull-downs; proteins with log₂ fold change ≥1 and P < 0.05 are highlighted. **(H)** Validation of FD3-ACDC interaction via GST pull-down and immunoblotting confirms association with full-length FD3.

To characterize the molecular interactions mediated by the FD3 ACDC domain we purified a recombinant GST-tagged FD3 ACDC protein (Fig. 3F) and performed pull-down assays coupled with LC-MS/MS analysis of gametocyte lysates from the *6HA::ap2-o3; 4Myc::fd3 parasites*. The FD3 ACDC pull-down recovered 481 significantly enriched proteins (Table S2). For the FD3 ACDC domain, the most strongly enriched protein was FD3, and AP2-O3 was not identified (Fig. 3G). Notably, the FD3 ACDC-enriched set also contained chromatin and transcription-associated proteins (Table S2). The homotypic interaction between the endogenous full-length FD3 and the shorter GST::ACDC domain bait was confirmed by immunoblotting (Fig. 3H). The failure of the FD3 ACDC bait to recover AP2-O3 suggests that the AP2-O3-FD3 interface may involve FD3 regions outside the isolated ACDC domain, that the association may be indirect or bridged by other factors, or that the FD3 ACDC domain preferentially engages in FD3 dimer under these conditions.

### ACDC deletion preserves selected female markers but alters a subset of asexual-stage transcripts in gametocytes

We performed RNA-seq on WT, *Δfd3*, and *fd3Δacdc* parasites to investigate how ACDC-mediated FD3 function influences gametocyte transcriptional programs (Fig. 4A). Genes showing differential abundance (hereafter referred to as differentially expressed genes, DEGs) were identified using DESeq2, with thresholds of FDR ≤ 0.05 and |log₂FC| ≥ 1. Compared with WT parasites, *fd3Δacdc* mutants exhibited 165 lower abundance and 187 higher abundance genes, whereas FD3 deletion resulted in a substantially broader transcriptional response, with 777 genes showing lower abundance and 402 showing higher abundance (Fig. 4B, C). We excluded the *pir* (*yir*) and *fam* multigene families from subsequent analyses because their expression is strongly influenced by stochastic epigenetic variation [23, 24]. After filtering, *fd3Δacdc* retained 58 lower-abundance and 169 higher-abundance DEGs. In comparison, *Δfd3* retained 754 genes with lower abundance and 248 genes with higher abundance (Fig. 4D). Only 29 lower-abundance and 28 higher-abundance genes overlapped between the two mutants (Fig. 4E, F). The substantially smaller DEG set in *fd3Δacdc* than in *Δfd3* indicates that the ACDC domain does not contribute significantly to the transcriptional activity of FD3 in gametocytes.

**Figure 4.**
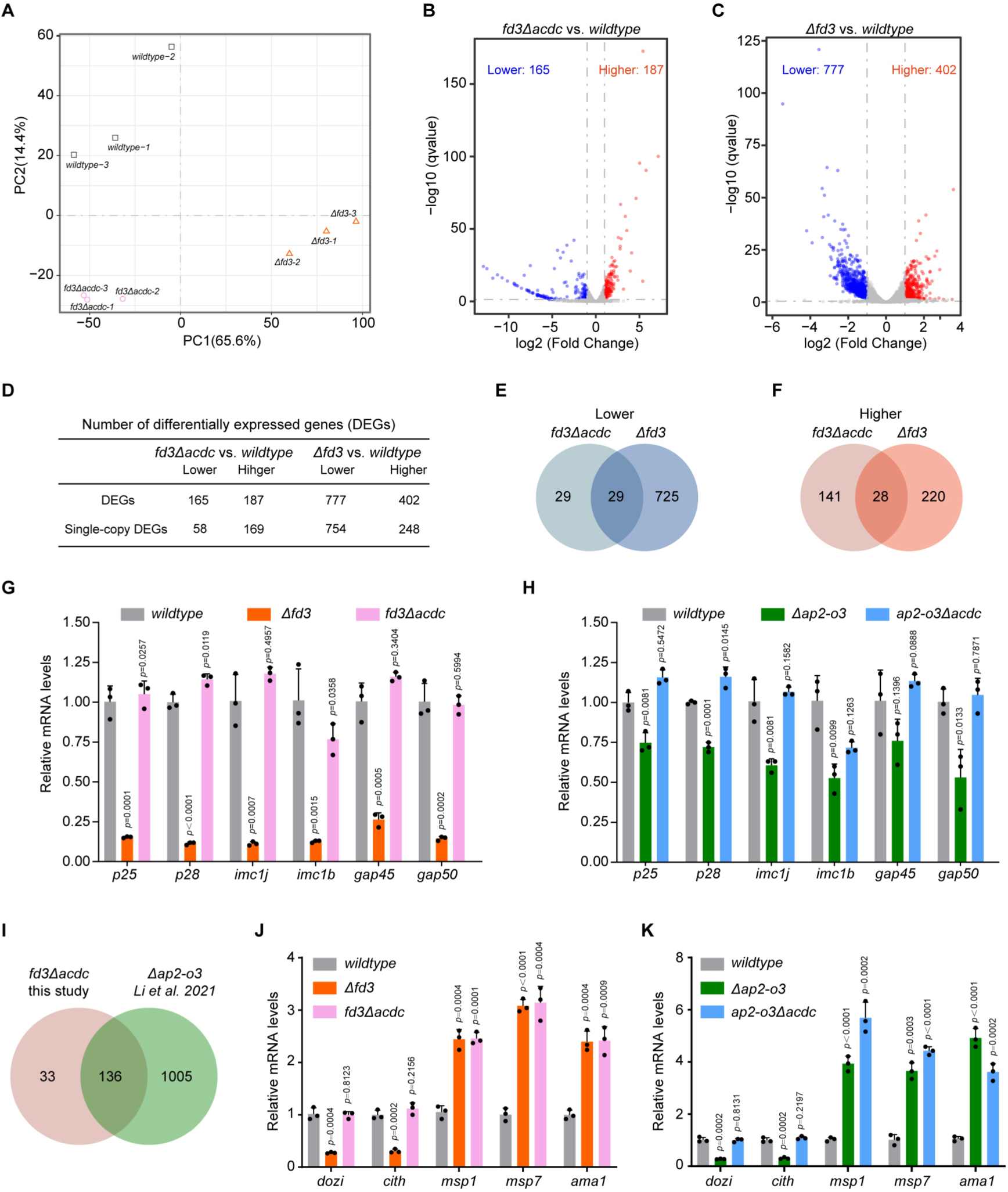
ACDC deletion preserves selected female gametocyte markers but alters a subset of asexual-stage transcripts in gametocytes. **(A)** Principal component analysis of RNA-seq profiles from WT, *Δfd3* and *fd3Δacdc* mixed gametocytes. **(B, C)** Volcano plots showing differentially abundant transcripts in *fd3Δacdc* and *Δfd3* parasites, respectively, relative to WT. Transcripts meeting the significance thresholds of |log₂ fold change| ≥ 1 and FDR ≤ 0.05 are shown in red for increased abundance and blue for decreased abundance. **(D)** Numbers of significantly differentially abundant transcripts in *fd3Δacdc* and *Δfd3* parasites after exclusion of the *pir*/*yir* and *fam* multigene families. **(E, F)** Venn diagrams showing overlap of transcripts with lower abundance (E) and higher abundance (F) between *fd3Δacdc* and *Δfd3* parasites. **(G, H)** qRT–PCR validation of selected female-specific gametocyte genes in *Δfd3* and *fd3Δacdc* parasites (G) and *Δap2-o3* and *ap2-o3Δacdc* parasites (H). **(I)** Venn diagram showing the overlap of genes with higher abundance in *fd3Δacdc* mutants (this study) and *Δap2-o3* female gametocytes (from *Li et al.*, 2021). **(J, K)** qRT–PCR analysis of selected female gametocyte genes (*dozi* and *cith*) and asexual-stage genes (*msp1*, *msp7* and *ama1*) in *Δfd3* and *fd3Δacdc* parasites (J) and *Δap2-o3* and *ap2-o3Δacdc* parasites (K). Transcript abundance was normalized to 18S rRNA and expressed relative to WT controls. Data are presented as mean ± SD from three biological replicates. Statistical significance was determined using a two-tailed Student’s *t*-test.

To further assess whether the ACDC domain is required for FD3-mediated maintenance of female gametocyte mRNA repertoire, we quantified transcripts of six well-characterized female-specific genes. Because the selected female-specific genes (*p25*, *p28*, *imc1j*, *imc1b*, *gap45*, and *gap50*) did not meet the transcriptome-wide differential-abundance thresholds in the *fd3Δacdc* RNA-seq analysis, we assessed them separately by targeted qRT-PCR to test whether ACDC deletion altered their abundance. In *Δfd3* mutants, all six transcripts were reduced to approximately 20% of WT levels, while *Δap2-o3* parasites showed higher transcript levels (50–75% of WT). Transcript levels in *fd3Δacdc* and *ap2-o3Δacdc* mutants were comparable to those in WT (Fig. 4G, H) suggesting that ACDC deletion did not impact the expression of these female gametocyte markers.

We next examined the molecular basis of the partial developmental defect in ACDC-deletion mutants. RNA-seq analysis revealed that transcript levels of several ookinete development genes were not substantially reduced in *fd3Δacdc* mutants. Given that AP2-O3 and FD3 function in a transcriptional repressor complex in female gametocytes [21], we compared the DEGs that were more abundant in *fd3Δacdc* mutants with those identified in *Δap2-o3* purified female gametocytes [19]. More than 80% (136/169) of genes with higher abundance in the FD3 ACDC deletion mutant were also significantly higher in abundance following AP2-O3 loss (Fig. 4I). We further analyzed the expression of representative asexual-stage genes in *Δfd3* and *fd3Δacdc* mutants by qRT-PCR. Transcripts of *msp1*, *msp7*, and *ama1* were significantly elevated in both mutants relative to WT in gametocytes. In contrast, *dozi* and *cith* transcripts were reduced in *Δfd3* parasites but remained largely unchanged in *fd3Δacdc* mutants (Fig. 4J). In addition, asexual-stage gene expression was also elevated in *Δap2-o3* and *ap2-o3Δacdc* mutants (Fig. 4K). Taken together, these data show that deletion of the FD3 ACDC domain is associated with increased abundance of a set of asexual-stage transcripts and reduced ookinete conversion, while the female gametocyte markers examined remain largely unchanged. The overlap between transcripts increased after FD3 ACDC deletion and those increased following AP2-O3 ACDC loss is consistent with related regulatory outcomes (Fig. 5).

**Figure 5.**
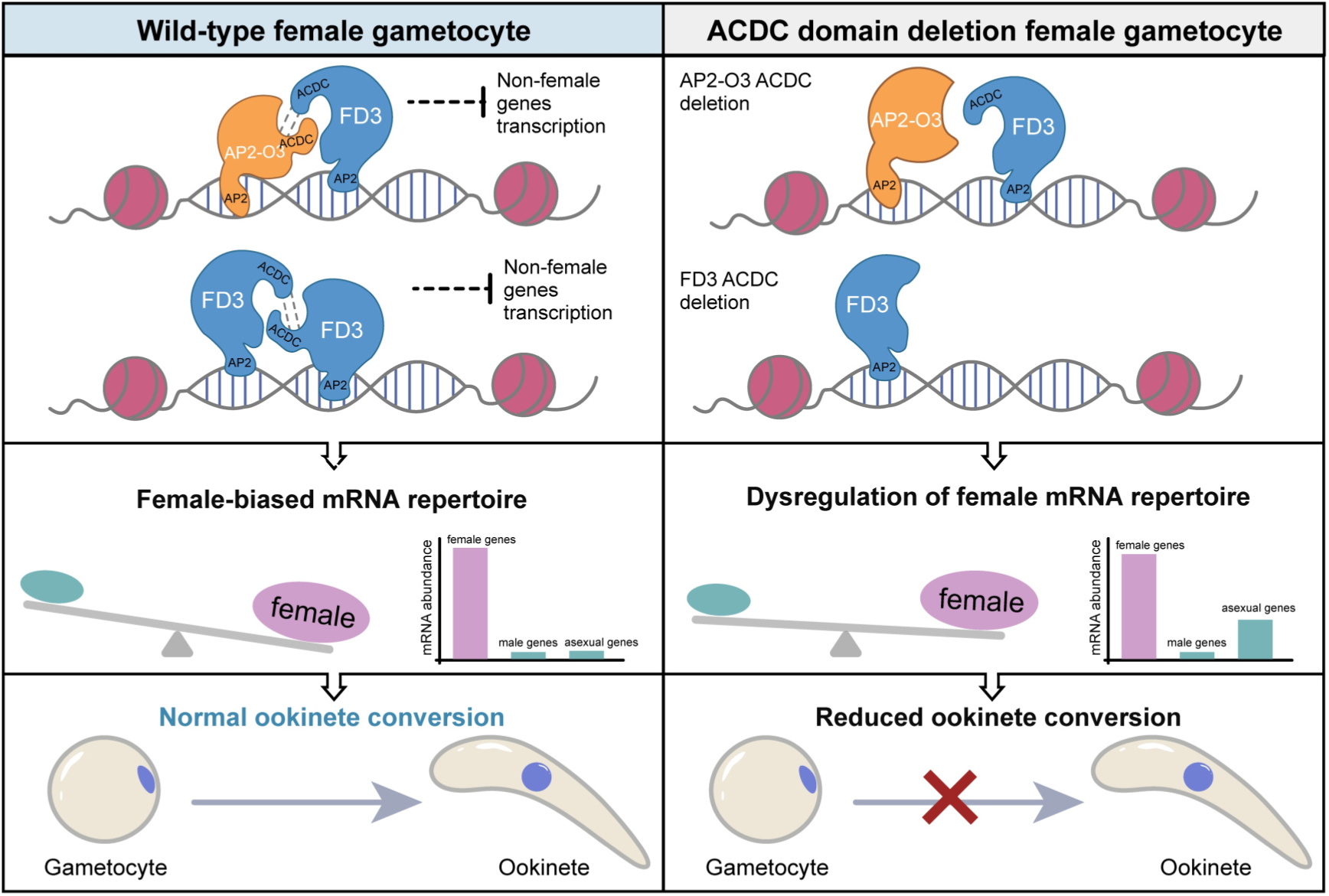
Proposed model of AP2-O3 and FD3 ACDC-dependent protein interactions and developmental regulation. In female gametocytes, AP2-O3 and FD3 are associated by co-immunoprecipitation, and the isolated AP2-O3 ACDC domain enriches FD3 in lysate-based pull-down assays. The FD3 ACDC domain preferentially recovers full-length FD3, consistent with possible self-association or an indirect association. The identities, directness, and stoichiometry of the relevant protein interactions remain unresolved. Deletion of either ACDC domain leaves the tested protein abundance, nuclear localization, and female gametocyte markers largely unchanged, but is associated with increased abundance of selected asexual-stage transcripts and reduced mature ookinete conversion.

### The *Pf*FD3 ACDC domain is dispensable for gametocyte maturation, female-specific expression, and nuclear localization

Given the essential contribution of the FD3 ACDC domain to transmission-stage development in *P. yoelii*, we next investigated whether this structural and functional requirement is conserved in the human malaria parasite *Plasmodium falciparum*. The *Pf*FD3 ortholog (PF3D7_1319600) contains a conserved C-terminal ACDC domain, and we generated a *Pf*FD3ΔACDC truncation line together with a matched full-length *Pf*FD3 control line (*Pf*FD3dd) using homologous recombination-based genome editing (Fig. S3A, B). Since *Pf*FD3 transcription is restricted to the sexual stages, we conducted IFAs on mixed asexual blood stages of the *Pf*FD3dd control line. No detectable *Pf*FD3 expression was observed in asexual parasites, and the *Pf*FD3ΔACDC truncation line exhibited normal asexual-stage development. This recapitulates work from both *P. berghei* and *P. yoelii* rodent systems (this study), reporting that FD3 is dispensable during asexual growth and is expressed only in gametocytes [20, 21]. We next assessed whether the *Pf*FD3 ACDC domain contributes to gametocyte development and maturation. Following induction of gametocytogenesis, the developmental progression of *Pf*FD3dd and *Pf*FD3ΔACDC parasites was monitored throughout gametocyte maturation. Growth curves normalized to day 2 post-induction parasitemia revealed no significant developmental delay or growth defect in the absence of the ACDC domain (Fig. S4). Detailed morphological staging from stage I to stage V further demonstrated that *Pf*FD3ΔACDC parasites progressed through all stages of gametocytogenesis without detectable developmental arrest, morphological abnormalities, or stage-specific defects compared with *Pf*FD3dd controls (Fig. 6A). To determine whether ACDC deletion affects the sex-specific expression pattern or nuclear localization of *Pf*FD3, we performed immunofluorescence assays using DDD and CITH as male- and female-specific markers, respectively [25, 26]. In both *Pf*FD3dd and *Pf*FD3ΔACDC parasites, *Pf*FD3 signals were detected exclusively in the nuclei of CITH-positive female gametocytes and were absent from DDD-positive male gametocytes. Importantly, removal of the ACDC domain did not alter female-specific expression or nuclear localization of *Pf*FD3 (Fig. 6B). Therefore, *Pf*FD3 is restricted to female gametocytes and does not require the ACDC domain for the production of mature gametocytes.

**Figure 6.**
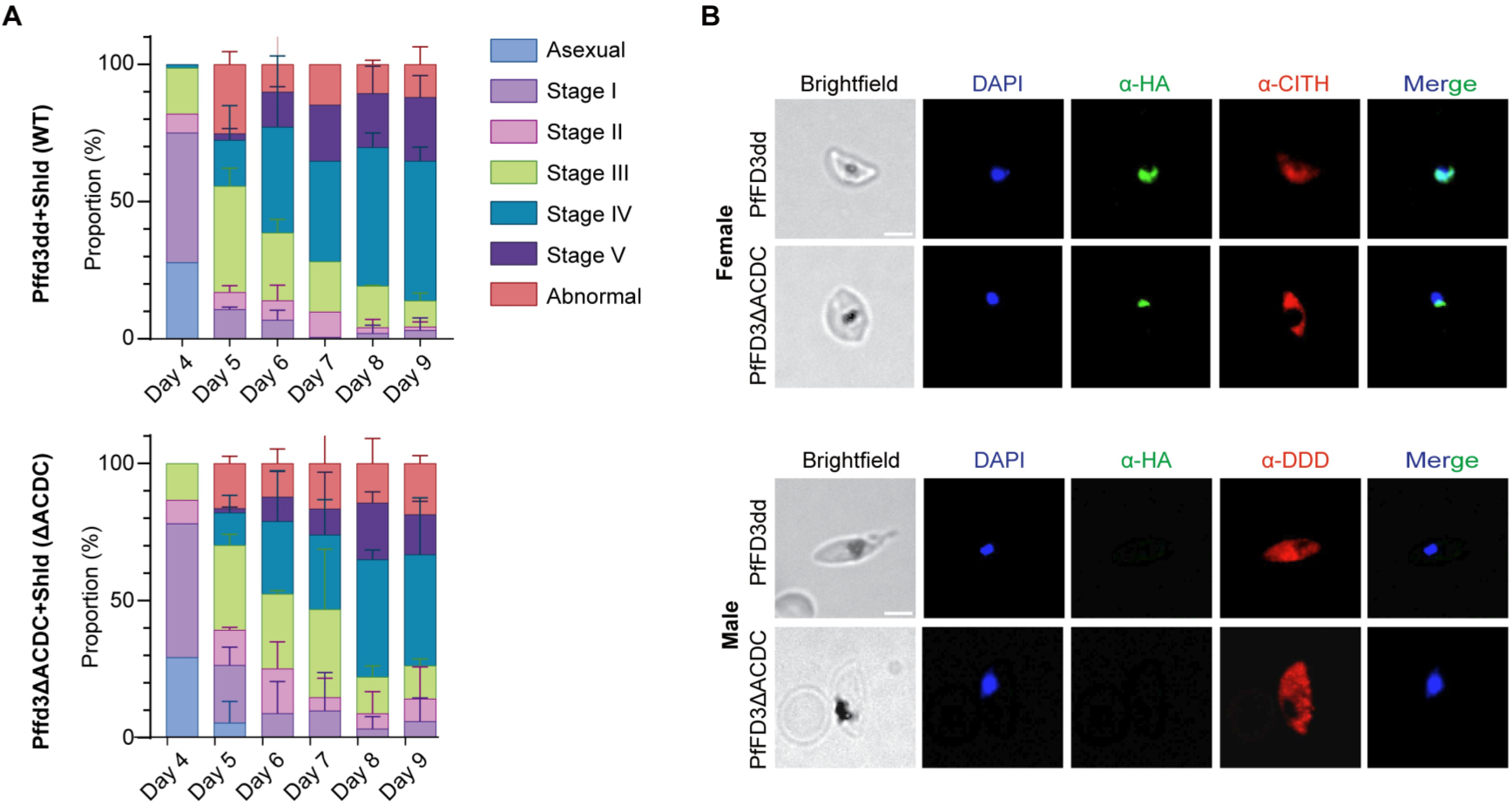
Subcellular localization of *P. falciparum* FD3 in the presence or absence of ACDC, and stage distribution of gametocytes during gametocyte development. (A) Staging of PfFD3dd and PfFD3ΔACDC parasites throughout gametocyte development. For each day, 50-200 gametocytes were counted. Counts are based on N=2 biological replicates. **(B)** Immunofluorescence microscopy of female and male PfFD3dd and PfFD3ΔACDC Stage III (day 7 post-induction) gametocytes. Parasites were stained with anti-HA to mark FD3 (green) and rabbit polyclonal antibodies against either PyCITH or PyDDD (red) as female and male markers, respectively. DNA was stained with DAPI (blue). Colocalization of nuclei and FD3 is shown in cyan in the merged image. Scale bars = 4 μm.

### Structure-guided virtual screening of conserved ACDC domains prioritizes candidate ligands for PfAP2-O3 and PfFD3

ACDC domains lack detectable human homologs, making them attractive candidates for structure-guided compound screening. We first compared the ACDC domains of *P. yoelii* and *P. falciparum* at both the sequence and structural levels. The AP2-O3 ACDC domains share 58% sequence identity and 75% sequence similarity between the two species, while the FD3 ACDC domains were more conserved, with 80% sequence identity and 90% sequence similarity (Fig. S5A, B). Structural superposition further revealed conserved predicted three-dimensional architectures, with RMSDs of 0.631 Å and 0.516 Å for AP2-O3 and FD3, respectively (Fig. S5C, D). We used Protenix to predict the three-dimensional structures of the *P. falciparum* AP2-O3 and FD3 ACDC domains for subsequent docking analyses [27]. Both predicted structures adopted compact four-helix bundles, consistent with the experimentally determined ACDC fold [14]. Putative ligand-binding cavities identified using CB-DOCK2 differed in their overall architectures: *Pf*AP2-O3-ACDC contained a relatively open, surface-exposed hydrophobic cavity (Fig. S5E), while *Pf*FD3-ACDC contained a more enclosed pocket within the helical core (Fig. S5F). These predicted cavities were subsequently used for independent structure-based virtual screening of the compound library.

The screening library consisted of 283 compounds: 276 dual-stage antimalarial compounds from the Tres Cantos Antimalarial Set (TCAMS), which have been evaluated in functional gametocyte assays [28, 29], plus seven previously reported small molecules targeting the ACDC domains of *Pf*AP2-I, *Pv*AP2-I, or *Pf*AP2-O5 from [14]. The 20 top-ranked compounds for the *Pf*AP2-O3 ACDC domain had predicted AutoDock Vina scores from −8.844 to −7.275 kcal/mol. The highest three were TCMDC-124223, TCMDC-139105, and TCMDC-133229 (Table 1). For the *Pf*FD3 ACDC domain, the top 20 compounds scored between −8.957 and −8.155 kcal/mol; the top three were TCMDC-134543, TCMDC-132053, and TCMDC-125432 (Table 2). Seven compounds, TCMDC-124223, TCMDC-124284, TCMDC-132053, TCMDC-132580, TCMDC-134543, TCMDC-137354, and TCMDC-138845, appeared in the top-20 lists for both domains, showing some overlap in compounds favored by the predicted binding cavities. However, their docking rankings varied between the two targets.

**Table 1:**
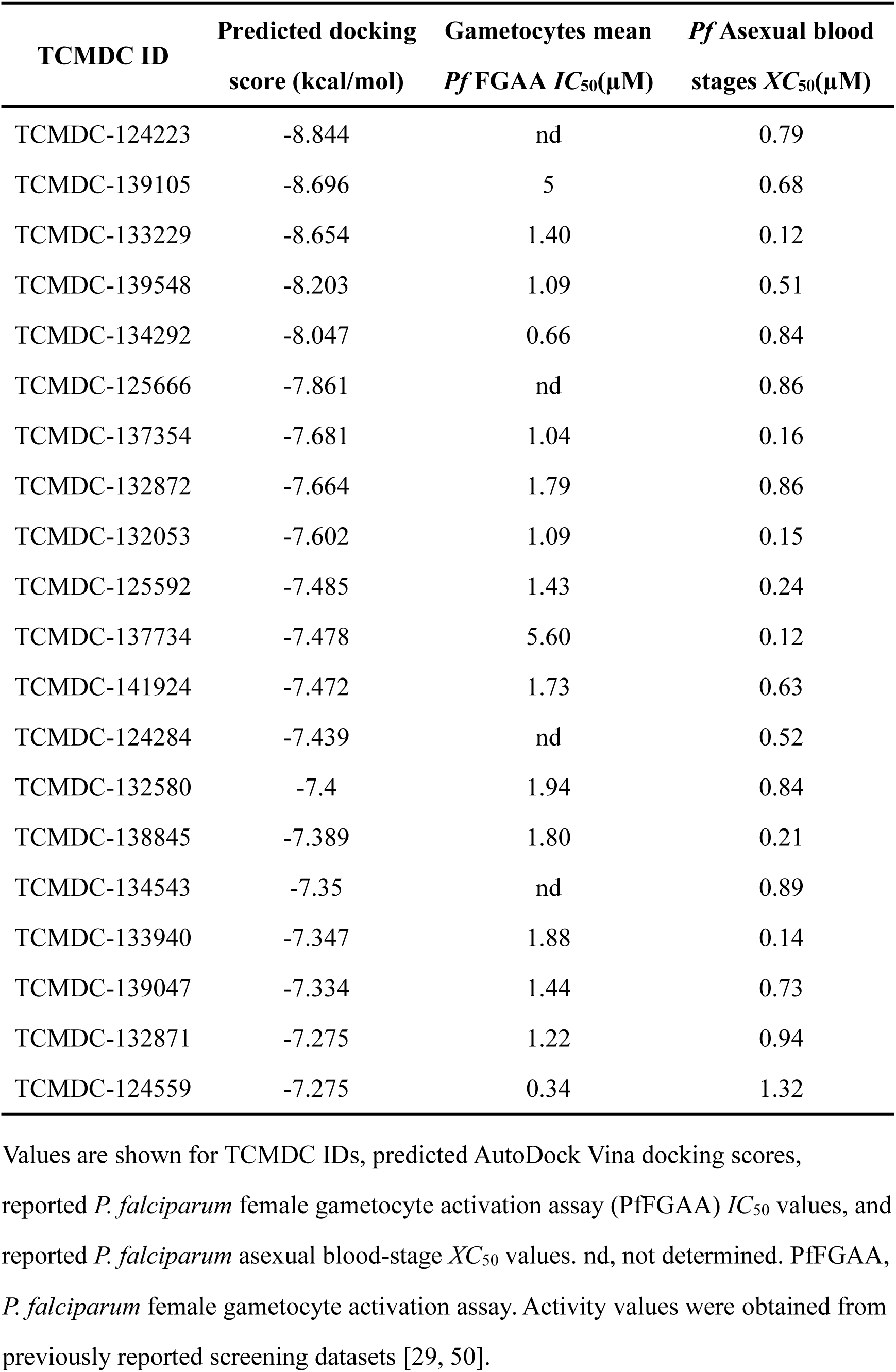
Top 20 compounds ranked by predicted AutoDock Vina docking score against the PfAP2-O3 ACDC domain, with reported *P. falciparum* inhibitory activities

**Table 2:** Top 20 compounds ranked by predicted AutoDock Vina docking score against the PfFD3 ACDC domain, with reported *P. falciparum* inhibitory activities

| **TCMDC ID** | **Predicted docking score (kcal/mol)** | **Gametocytes mean *Pf* FGAA *IC*<sub>50</sub> (µM)** | ***Pf* Asexual blood stages *XC*<sub>50</sub> (µM)** |
|---|---|---|---|
| TCMDC-134543 | -8.957 | nd | 0.89 |
| TCMDC-132053 | -8.909 | 1.09 | 0.15 |
| TCMDC-125432 | -8.82 | 1.07 | 0.83 |
| TCMDC-124284 | -8.777 | nd | 0.52 |
| TCMDC-137354 | -8.721 | 1.04 | 0.16 |
| TCMDC-132243 | -8.7 | 1.83 | 0.70 |
| TCMDC-140775 | -8.666 | 2.00 | 0.71 |
| TCMDC-138845 | -8.561 | 1.80 | 0.21 |
| TCMDC-124223 | -8.493 | nd | 0.79 |
| TCMDC-124787 | -8.486 | 2.12 | 0.75 |
| TCMDC-134436 | -8.444 | 1.08 | 0.99 |
| TCMDC-140112 | -8.422 | 1.38 | 0.16 |
| TCMDC-123754 | -8.417 | 1.16 | 0.44 |
| TCMDC-137820 | -8.404 | 1.06 | 0.003 |
| TCMDC-132580 | -8.295 | 1.94 | 0.84 |
| TCMDC-132811 | -8.281 | 0.83 | 0.26 |
| TCMDC-137813 | -8.181 | 2.01 | 0.05 |
| TCMDC-136658 | -8.168 | 1.46 | 0.45 |
| TCMDC-134444 | -8.165 | 1.32 | 0.90 |
| TCMDC-133882 | -8.155 | 1.30 | 0.15 |

Values are shown for TCMDC IDs, predicted AutoDock Vina docking scores, reported *P. falciparum* female gametocyte activation assay (PfFGAA) *IC*<sub>50</sub> values, and reported *P. falciparum* asexual blood-stage *XC*<sub>50</sub> values. nd, not determined. PfFGAA, *P. falciparum* female gametocyte activation assay. Activity values were obtained from previously reported screening datasets.

We then predicted the binding poses and protein–ligand interaction profiles for the top three compounds associated with each ACDC domain. For the *Pf*AP2-O3 ACDC domain, TCMDC-124223 was predicted to establish hydrophobic contacts with Ile1098, Leu1106, Phe1110, Ile1117, Tyr1120, Phe1140, Asn1147, Leu1149, and Pro1150, along with a hydrogen bond involving Tyr1120 (Fig. 7A). TCMDC-139105 showed a similar but partially overlapping interaction pattern, with hydrophobic contacts involving Leu1095, Ile1098, Leu1106, Val1113, Ile1117, Tyr1120, and Phe1140, as well as hydrogen bonds with Val1144 and Asn1147 (Fig. 7B). TCMDC-133229 was predicted to interact hydrophobically with Leu1106, Phe1110, Val1113, Leu1149, and Pro1150 (Fig. 7C). Notably, Tyr1120 was consistently identified as key interaction residue across these complexes (Fig. 7D). In the *Pf*FD3 ACDC domain, TCMDC-134543 was predicted to form hydrophobic contacts with Ile1608, Leu1614, Gln1618, Tyr1621, and Ile1622, along with a π-stacking interaction with Tyr1621 and a halogen bond involving Tyr1578 (Fig. 7E). TCMDC-132053 showed predicted hydrophobic contacts with Tyr1578, Gln1618, and Tyr1621, as well as two π-stacking interactions involving Tyr1621 (Fig. 7F). TCMDC-125432 was predicted to form hydrophobic contacts with Tyr1578, Lys1581, Gln1618, Tyr1621, and Ile1622, plus a π-stacking interaction with Tyr1621 (Fig. 7G). Gln1618 and Tyr1621 were consistently identified across all three *Pf*FD3 complexes, while Tyr1578 and Ile1622 were also observed in multiple complexes (Fig. 7H). The top-ranked compounds mainly exhibited hydrophobic interactions, with some also forming hydrogen bonds in the *Pf*AP2-O3 ACDC complexes and frequently engaging in π-stacking in the *Pf*FD3 complexes. The highest-ranked compounds for each domain, TCMDC-124223 for PfAP2-O3 and TCMDC-134543 for PfFD3, were then chosen for further molecular dynamics analysis.

**Figure 7.**
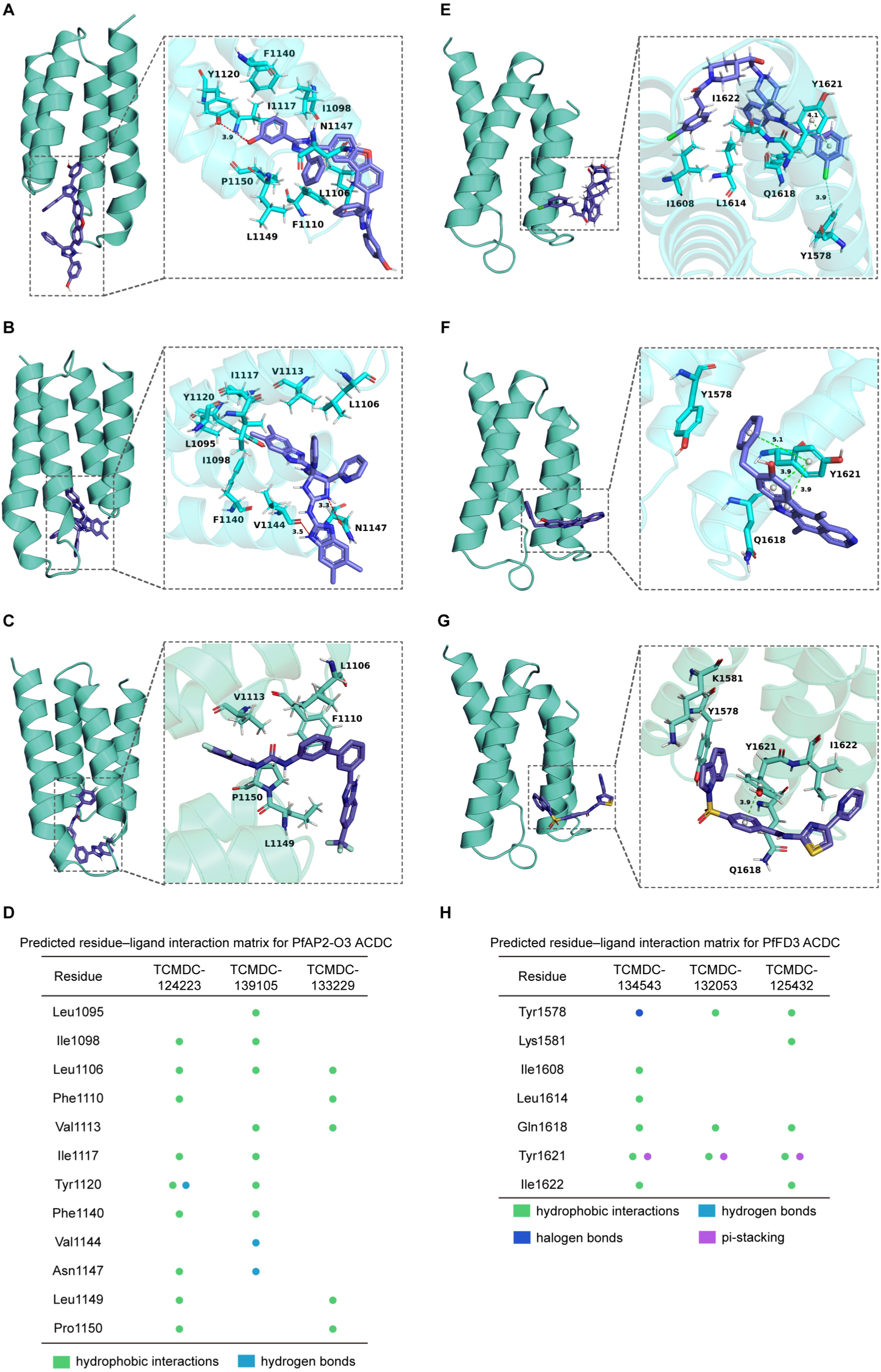
Predicted binding modes and residue-level interaction profiles of top-ranked compounds within the PfAP2-O3 and PfFD3 ACDC domains. **(A-C)** Predicted binding poses of the three highest-ranked compounds TCMDC-124223, TCMDC-139105, and TCMDC-133229 within the PfAP2-O3 ACDC domain. **(D)** Matrix summarizing the predicted residue-ligand interactions across the three PfAP2-O3 ACDC complexes. **(E-G)** Predicted binding poses of TCMDC-134543, TCMDC-132053, and TCMDC-125432 within the PfFD3 ACDC domain. **(H)** Matrix summarizing the predicted residue-ligand interactions across the three PfFD3 ACDC complexes. Hydrophobic interactions, hydrogen bonds, halogen bonds, and π-stacking interactions are indicated by green, cyan, blue, and purple symbols, respectively. Interactions and geometric distances were assigned by PLIP using default criteria.

### Molecular dynamics supports the persistence of the predicted top-ranked ACDC–ligand complexes

To further characterize the conformational behavior of the predicted ACDC–ligand complexes, the *Pf*AP2-O3 ACDC–TCMDC-124223 and *Pf*FD3 ACDC–TCMDC-134543 systems were subjected to 350-ns all-atom molecular dynamics simulations. Protein backbone RMSD profiles were analyzed for both ligand-free and ligand-bound systems. In the *Pf*AP2-O3 ACDC–TCMDC-124223 complex, the RMSD initially adjusted before stabilizing later in the simulation. The ligand-bound system reached a stable RMSD faster and showed fewer fluctuations after stabilization compared to the ligand-free system (Fig. 8A). Snapshots revealed that TCMDC-124223 moved and changed conformation within the predicted binding cavity during the early simulation phase but remained bound without dissociation over the 350-ns trajectory (Fig. 8B). Residue RMSF analysis indicated that fluctuations were spread across the *Pf*AP2-O3 ACDC domain, with higher values at the N- and C-termini than in the core structured region. Binding of TCMDC-124223 reduced RMSF at multiple residues, suggesting decreased local flexibility in the ligand-bound state (Fig. 8C). Furthermore, an initial transient increase in the radius of gyration occurred early in the ligand-bound simulation, corresponding to the conformational adjustment of the ligand within the binding cavity, but afterward it stabilized around 1.25 nm (Fig. 8D).

**Figure 8.**
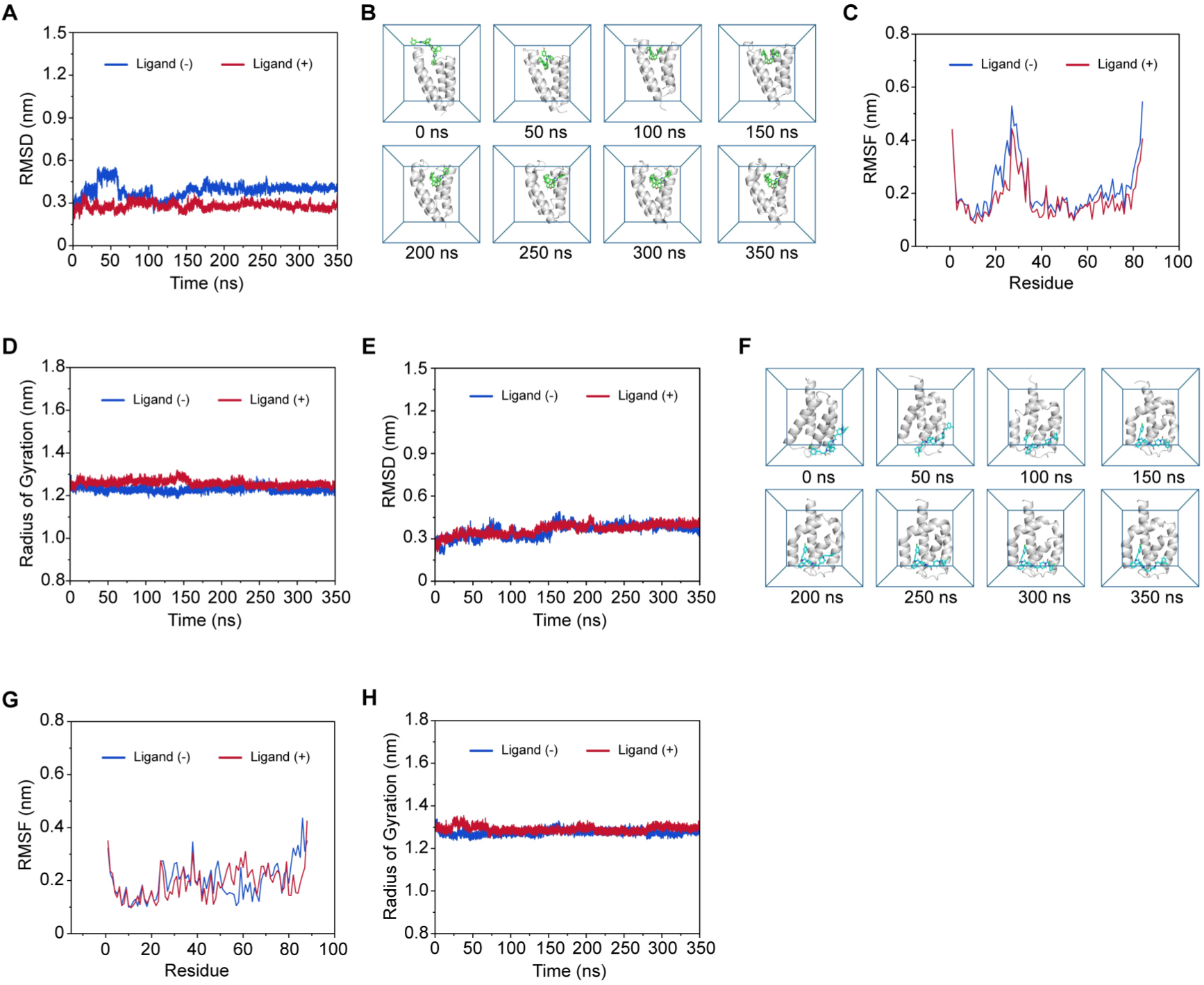
Molecular dynamics analysis of the conformational stability of the PfAP2-O3 and PfFD3 ACDC domains in the presence and absence of their prioritized ligands. **(A, E)** Protein RMSD profiles of the PfAP2-O3 ACDC and PfFD3 ACDC domains, respectively, in ligand-free and ligand-bound systems during 350-ns molecular dynamics simulations. **(B, F)** Representative snapshots of the PfAP2-O3 ACDC–TCMDC-124223 and PfFD3 ACDC–TCMDC-134543 trajectories at the indicated time points. **(C, G)** Residue-level RMSF profiles of the corresponding ACDC domains in ligand-free and ligand-bound systems. **(D, H)** Radius of gyration of the corresponding ACDC domains during the simulations.

The *Pf*FD3 ACDC–TCMDC-134543 complex showed a similar trajectory pattern. The backbone RMSD reflected an initial adjustment followed by convergence, with the ligand-bound system stabilizing faster and displaying fewer fluctuations in the later stages than the ligand-free system (Fig. 8E). Snapshots revealed that TCMDC-134543 initially moved and changed conformation within the predicted binding cavity during early simulation stages, after which it remained within the cavity without dissociation over 350 ns (Fig. 8F). RMSF analysis indicated that fluctuations varied by residue, mostly higher at the N- and C-termini. Many residues had lower RMSF values when bound to the ligand, suggesting decreased local flexibility upon TCMDC-134543 binding (Fig. 8G). During the first ∼75 ns, the Rg showed some variation as TCMDC-134543 adjusted within the cavity; afterward, the Rg stayed around 1.30 nm, indicating stable overall domain dimensions throughout the rest of the simulation (Fig. 8H). In both complexes, ligand binding led to faster RMSD convergence, fewer fluctuations at various residues, and the ligand remaining within the predicted binding cavity during later stages of the trajectory. These findings support sustained ligand binding and decreased conformational fluctuations of the protein in the ligand-bound states.

The interaction profiles derived from the molecular dynamics trajectories further support the idea that many of the contacts predicted by docking were preserved throughout the simulations. In the *Pf*AP2-O3 ACDC–TCMDC-124223 complex, residues such as Phe1110, Tyr1120, Phe1140, and Pro1150 repeatedly interacted with the ligand, mainly through hydrophobic and van der Waals forces. In the later stages of the trajectory, about two to three hydrogen bonds were maintained between the protein and ligand, with Tyr1120 showing the highest hydrogen-bond occupancy at 31.7% (Fig. S6A, B). Similarly, in the *Pf*FD3 ACDC–TCMDC-134543 complex, residues including Tyr1578, Leu1614, Thr1617, Gln1618, Tyr1621, Ile1622, Val1625, and Leu1626 engaged consistently with the ligand. Several residues from the docking models, notably Tyr1578, Leu1614, Gln1618, Tyr1621, and Ile1622, remained involved during the simulation. About one to two hydrogen bonds persisted in the later part of the simulation (Fig. S6D, E).

MM/GBSA analysis was performed using frames from 340–350 ns for the PfAP2-O3 ACDC–TCMDC-124223 complex and from 90–350 ns for the *Pf*FD3 ACDC– TCMDC-134543 complex. The effective binding free energies (ΔG_MM/GBSA) were −68.2 and −57.1 kcal/mol, respectively. In both cases, van der Waals interactions provided the main favorable energy contribution (−63.4 and −52.6 kcal/mol), while electrostatic contributions were comparatively minor (−1.2 and −0.4 kcal/mol); the polar solvation terms (GB) were +4.1 and +2.6 kcal/mol, and the nonpolar solvation terms (SASA) were −7.6 and −6.8 kcal/mol. These values should be regarded as relative estimates that support favorable, predominantly hydrophobic binding, rather than absolute affinities (Fig. S6C, F). These results agree with the dominance of hydrophobic and van der Waals contacts seen in both docking and trajectory analyses.

Collectively, our structure-based screening and molecular dynamics analyses prioritize TCMDC-124223 and TCMDC-134543 as candidate ligands for the PfAP2-O3 and PfFD3 ACDC domains, respectively. Deletion of the ACDC domain in *Py* did not produce an asexual blood-stage growth defect, consistent with the stage-restricted functions of AP2-O3 and FD3. We observed partial inhibition of gametocyte-to-ookinete conversion upon ACDC deletion, demonstrating a functional requirement for ACDC domains during parasite transmission-stage development and supporting ACDC as a candidate target for transmission-blocking interventions. High sequence and structural conservation, especially for FD3 ACDC domains between *Py* and *Pf*, suggests the potential for cross-species ligand binding, though equivalent binding affinity and in vivo activity await validation.

## Discussion

ApiAP2 proteins constitute the major family of transcriptional regulators in *Plasmodium* and are involved in stage-specific gene expression programs throughout the parasite’s life cycle [6, 8–12]. How a relatively small number of ApiAP2 factors generate diverse, context-specific regulatory outputs beyond sequence-specific DNA recognition remains poorly understood. In this study, we provide evidence that the apicomplexan-specific ACDC domain is involved in protein interactions involving ApiAP2 factors. In *P. yoelii* gametocytes, AP2-O3 and FD3 co-immunoprecipitated, the AP2-O3 ACDC bait enriched FD3, and the FD3 ACDC bait preferentially recovered full-length FD3. These findings support an association between AP2-O3 and FD3 and are consistent with possible FD3 self-association or an indirect association involving full-length FD3. However, our experiments do not establish direct binding between these proteins or define a higher order protein complex. Functionally, ACDC deletion partially impaired the transition from gametocytes to ookinetes without substantially altering the female gametocyte markers. These findings broaden previous observations that AP2-O3 and FD3 function within a transcriptional repressor complex [21] and suggest that ACDC-mediated protein interactions contribute to the complex regulatory activity during transmission-stage development. A role in complex formation has been demonstrated for the ACDC domain of *Pf*AP2-I [30], and ACDC-containing ApiAP2 factors in *Toxoplasma gondii* likewise operate within multimeric regulatory complexes [31, 32]. These findings suggest that ACDC domains may serve as structural modules that facilitate protein associations involving ApiAP2 factors.

The truncated ACDC-deleted AP2-O3 and FD3 ApiAP2 proteins were still detectable and localized to the nucleus, and transcription within female gametocytes remained largely intact. Conversely, complete disruption of AP2-O3 or FD3 caused far more extensive transcriptional and developmental disturbances [19–21]. Thus, ACDC deletion does not remove the core regulatory functions required to establish female gametocyte identity. Instead, it impairs specific aspects of their regulatory activity in female gametocytes. Since AP2-O3 mainly acts as a transcriptional repressor and FD3 promotes female-specific transcription, different experimental approaches are needed to study these factors. In particular, the effects of AP2-O3 loss are best assessed in purified female gametocytes, where increased expression of male- or asexual-related genes clearly indicates a loss of repression. In mixed populations, such interpretations are limited because changes in male transcripts may also reflect altered cell composition. In contrast, the role of FD3 can be more directly examined in mixed populations by monitoring the decrease in female-specific transcripts, which suggests impaired transcriptional activation or maintenance. Therefore, our RNA-seq analysis focused on the FD3 ACDC deletion, while AP2-O3 effects were assessed through targeted expression studies and by comparing with previously published AP2-O3 loss-of-function transcriptomes. In line with this experimental reasoning, deleting either AP2-O3 or FD3 ACDC resulted in elevated levels of specific asexual-stage transcripts. Furthermore, many genes that showed increased transcript levels after FD3 ACDC deletion also exhibited higher levels after AP2-O3 loss. These shared transcriptional alterations, together with the partial ookinete-conversion phenotype, suggest that ACDC-dependent protein associations may contribute to transcriptional regulation during transmission-stage development. However, the current data do not establish a direct mechanistic link between a defined protein assembly and these transcriptional changes. ACDC-dependent associations may contribute to maintaining the balance between female-specific and non-female gene expression programs, a possibility that warrants further investigation.

Deleting the FD3 ACDC domain in *P. falciparum* showed no impact on asexual growth, female-specific expression, nuclear localization, or gametocyte maturation through stage V. These results are consistent with the *P. yoelii* findings, indicating that the ACDC domain is not essential for key aspects of gametocyte development across species. However, the *P. falciparum* analysis was limited to FD3 and did not evaluate the roles of AP2-O3 or stages after gametocyte transmission. It remains unclear whether ACDC-mediated interactions affect gametocyte activation, fertilization, ookinete formation, or mosquito-stage development. Additionally, differences in gametocyte biology between rodent and human malaria parasites suggest caution when applying *P. yoelii* findings to *P. falciparum* transmission [33–35]. Gaining a complete understanding of the function of these domains in *P. falciparum* transmission will necessitate studying more developmental stages and directly testing AP2-O3/FD3 co-association and its dependence on the ACDC domains.

The structure-based predictions provided a computational framework for prioritizing compounds targeting the two ACDC domains. While each domain produced different compound rankings, several compounds were favored by both binding cavities. Notably, four compounds—TCMDC-124223, TCMDC-124284, TCMDC-137354, and TCMDC-134543—were among the shared top candidates in this study and also ranked highly in prior structure-based screenings against the ACDC domains of *Pf*AP2-I, *Pv*AP2-I, and *Pf*AP2-O5 [14]. The repeated presence of these compounds in ACDC domains corresponds to the known structural conservation of a hydrophobic ligand-binding pocket identified in ACDC-containing ApiAP2 proteins [14]. In line with this observation, the interaction profiles predicted here were characterized by hydrophobic contacts, with multiple aromatic and polar residues consistently appearing across the highest-ranking complexes. Notably, Tyr1578 in *Pf*FD3 was previously identified as corresponding to Tyr626 in the *Pf*AP2-O5 ACDC domain, a residue involved in ligand binding within the conserved ACDC pocket [14]. These findings support the idea of a recurring structural environment compatible with ligand binding across multiple ACDC domains. They also imply that the ACDC domain might be a common, targetable module among various ApiAP2 regulators that operate at different stages of the parasite’s life cycle. This is especially significant because ApiAP2 proteins with the ACDC domain participate in various developmental processes, such as AP2-I-dependent regulation of erythrocyte invasion and functions associated with AP2-O3 and AP2-O5 during mosquito-stage development [8]. However, experimental confirmation awaits to determine whether certain compounds can bind to ACDC domains in parasites, interfere with their protein interactions, and lead to stage-specific or multi-stage antimalarial effects.

To our knowledge, this is the first study to define a functional role for ACDC domains in *Plasmodium* parasites. Our findings support a model in which the ACDC domain mediates protein interactions between ApiAP2 proteins. For AP2-O3 and FD3, the ACDC domains facilitate the formation of a regulatory complex that maintains a unique transcriptional repertoire for female gametocytes and ensures a smooth transition to ookinetes. The observed partial phenotypes after ACDC deletion suggest that while the domain affects the regulatory functions of these proteins, it does not fully control them. Furthermore, the conserved structure of ACDC and its capacity to connect with regulatory partners suggest that the organization of protein complexes adds an extra layer of specificity to ApiAP2 functions, beyond sequence-specific DNA binding. These structural features also lay the groundwork for future drug development efforts aimed at blocking transmission.

## Materials and Methods

### Ethics statement

All mouse experiments were conducted in strict accordance with the guidelines of the Institutional Animal Use and Ethics Committee of the University of South China (Hengyang, China).

### Animal and parasite usage

Female ICR mice (5-7 weeks old) were used for parasite propagation and monoclonal selection. The mice were obtained from Hunan SJA Laboratory Animal Co. Ltd. (Changsha, China, Animal Production License No. SCXK 2021-0002). All transgenic parasite lines were generated from the *Plasmodium yoelii* 17XNL strain using the CRISPR/Cas9 method [36]. A complete list of all parasite strains generated in this study is provided in Table S3.

### Plasmid construction for *P. yoelii* genome modification

Plasmid construction for *P. yoelii* genome modification was performed using the CRISPR/Cas9 system as described previously [36, 37], with modifications tailored for the pYCm vector. Briefly, 5’ and 3’ flanking sequences (300-800 bp each) flanking the intended insertion site were amplified as homology templates and cloned into pYCm. For gene tagging, a DNA fragment encoding 6HA or 4Myc was inserted in-frame between the homology arms; for domain deletion, the left and right homologous arms flanking the target domain were directly ligated into the corresponding restriction sites after PCR amplification. At least two sgRNAs per target were designed using EuPaGDT (http://grna.ctegd.uga.edu/). All primers and oligonucleotides used in this study are listed in Table S3.

### Plasmid construction for *P. falciparum* genome modification

To generate the PfFD3-GFP-HA-DD24-glmS knock-in construct, two homology fragments were amplified from *P. falciparum* 3D7 genomic DNA. A 981-bp fragment corresponding to the 3’ end of PfFD3 (excluding the stop codon) was amplified with primers IP47_FD3_KD_HR_Fw and IP48_FD3_KD_HR_Rv. For the ACDC-truncated variant, a 983-bp upstream fragment flanking the ACDC domain was amplified with primers IP49_FD3_ACDCtrun_HR_Fw and IP50_FD3_ACDCtrun_HR_Rv (primer sequences listed in Table S3). Genomic DNA was extracted using the DNeasy Blood & Tissue Kit (QIAGEN), and primers were designed with Primer3. PCR was performed with LA Taq Advantage Genomic LA Polymerase Mix (Takara). Amplified products were purified, cloned into the pGEM-T Easy vector (Promega) via AT cloning, and transformed into XL10-Gold chemically competent *E. coli*. Transformants were selected on LB-ampicillin plates supplemented with 0.1 mM IPTG and 20 µg/mL X-gal. Eight positive colonies were grown in LB-ampicillin liquid medium, and plasmids were extracted using the QIAprep Spin Miniprep Kit (QIAGEN). Clones were verified by Sanger sequencing and aligned against the *P. falciparum* 3D7 reference sequence in PlasmoDB. Sequence-confirmed inserts were excised from pGEM-T Easy with PstI and BglII and ligated into the p_GFPHADD24AbsDglmS vector backbone [38], yielding the final transfection plasmid pFD3-GFP-HA-FKBP2A-bsd-glmS. This construct comprises the PfFD3 homology region fused in-frame to a GFP-HA tag, the ddFKBP destabilization domain DD24, a 2A self-cleaving peptide, the blasticidin S deaminase resistance cassette, the glmS ribozyme, and the hDHFR cassette for WR99210 resistance.

### Plasmodium falciparum culture

Parasites were maintained in RPMI 1640 medium supplemented with 0.25% AlbuMAX, 25 mM HEPES, 0.2% sodium bicarbonate, 13.6 µg/mL hypoxanthine, and 50 µg/mL gentamicin at 3% hematocrit, and incubated at 37 °C under an atmosphere of 6% O₂ and 5% CO₂. Gametocyte conversion was induced by treating sorbitol-synchronized parasites with mFA medium, following previously established protocols [39, 40]. At the time of mFA induction, all cultures were supplemented with 0.5 µM Shld-1 (provided by the Goldberg laboratory, Washington University in St. Louis). Approximately 22 h post-induction, the medium was replaced with standard RPMI 1640. To prevent reinvasion by asexual parasites, heparin (Sigma, H4784) was added to the cultures on day 2 post-induction. Parasitemia and gametocyte staging were monitored daily by light microscopy using 100× oil-immersion objective lenses on Giemsa-stained blood smears.

### *P. yoelii* parasite transfection and gene modification

Schizonts were isolated from infected mice by 60% Nycodenz density gradient centrifugation, washed twice with RPMI-1640, and electroporated with 5 μg of circular plasmid DNA using a Nucleofector 2b Device (Lonza, Germany). The transfected parasites were immediately injected into naive mice via the tail vein. Pyrimethamine selection (6 mg/ml in drinking water) was initiated on the following day, and resistant parasites were typically observed approximately 7 days later. Single-clone parasites were obtained by limiting dilution in mice. Genomic DNA was extracted from the blood of infected mice, and PCR genotyping was performed using the gene-specific primers listed in Table S3.

### Parasite negative selection with 5-fluorocytosine

To remove the episomal pYCm plasmid, we performed negative selection with 5-fluorocytosine (5FC; Aladdin, cat# C123460). The 5FC was dissolved in water to a final concentration of 2.0 mg/mL and provided to mice infected with the modified monoclonal parasite line in dark drinking bottles. After approximately three days, most surviving parasites no longer carried the pYCm plasmid; these parasites were then inoculated into mice via tail vein injection for limiting dilution cloning. Seven days later, PCR genotyping was performed to determine whether the pYCm plasmid had been completely cleared from the parasite populations. To further confirm plasmid loss, parasites that had undergone negative selection were subjected to a pyrimethamine (6 mg/ml in drinking water) sensitivity test. Only those parasites successfully cured of the pYCm plasmid were used in subsequent rounds of genome editing.

### Gametocyte induction in mouse

To induce reticulocytosis and promote *Plasmodium* gametocyte formation, ICR mice were intraperitoneally injected with phenylhydrazine (80 µg/g body weight; Sangon Biotech, cat#A600705). Three days post-treatment, each mouse was infected with 5×10⁶ *Plasmodium* parasites via tail vein injection, with peak gametocytaemia occurring on day three post-infection. Thin blood smears were prepared, stained with Giemsa, and male and female gametocytes were enumerated under a microscope. Gametocytaemia was calculated as the percentage of gametocytes relative to the total number of infected erythrocytes. All experiments were performed in triplicate.

### *In vitro* ookinete culture

*In vitro* culture for ookinete development was prepared as described previously [19]. Mouse blood containing gametocytes was collected in heparin-containing tubes and immediately mixed with the ookinete culture medium. The medium consisted of RPMI 1640 supplemented with 25 mM HEPES, 10% fetal calf serum, and 100 µM XA, pH 8.0. The blood and medium were mixed at a 1:10 ratio and incubated at 22 °C for 20 hours, after which thin blood smears were prepared and stained with Giemsa for analysis. The ookinete conversion rate was calculated as the number of ookinetes (including both mature and immature forms) per 100 female gametocytes. The mature ookinete conversion rate was calculated as the number of mature ookinetes per 100 female gametocytes. All experiments were performed in triplicate.

### Protein extraction and Western blot

Parasite pellets were lysed in RIPA buffer (Epizyme, cat#PC101) containing a protease inhibitor cocktail (MedChemExpress, HY-K0010). After ultrasonication on ice, the lysate was incubated on ice for 30 min and then centrifuged at 12,000 × g for 10 min at 4 °C. The resulting supernatant was collected. The protein samples were mixed with SDS-PAGE loading buffer, heated at 95 °C for 10 min, resolved by SDS-PAGE (with separating gel concentrations adjusted according to the target protein molecular weight), and transferred onto PVDF membranes (Millipore, cat#IPVH00010). The membranes were blocked with 5% non-fat milk in 1 × TBST buffer at room temperature for 2 h, followed by incubation with primary antibodies overnight at 4 °C. After three washes with TBST, the membranes were incubated with horseradish peroxidase-conjugated goat anti-rabbit or anti-mouse secondary antibodies for 60 min at room temperature and then washed again with TBST. Finally, the signals were visualized using a high-sensitivity ECL chemiluminescence detection kit (Abbkine, cat#BMU102-CN).

### Immunofluorescence assay for *P. yoelii*

Immunofluorescence assay for *P. yoelii* was performed as described previously [19]. Briefly, purified parasites were gently mixed with 4% paraformaldehyde solution (Servicebio, cat#G1101) and transferred to a 24-well cell culture plate containing poly-L-lysine-coated coverslips (Solarbio, cat#YA0350). After incubation for 10 min, the plate was centrifuged at 300 × g for 10 min to immobilize the parasites onto the coverslips, followed by two washes with PBS. The parasites were then permeabilized with 0.1% Triton X-100 in PBS for 7 min at room temperature, washed, and blocked with 5% BSA in PBS for 2 h. Thereafter, the samples were incubated with primary antibodies diluted in 5% BSA/PBS overnight at 4 °C. After washing, the coverslips were incubated with fluorescently conjugated secondary antibodies for 2 h at room temperature, during which nuclei were counterstained with Hoechst 33342 (Thermo Fisher, cat#62249). Finally, the coverslips were mounted in 90% glycerol and sealed with nail polish. All images were acquired and processed using a Zeiss LSM 880 laser scanning confocal microscope under identical settings.

### Immunofluorescence assay for *P. falciparum*

Immunofluorescence assays were adapted from [41]. Briefly, gametocytes were purified via magnetic separation (MACS LS Columns, Miltenyl Biotec) and then fixed in 4% paraformaldehyde (Thermo 28908) and 0.0075% glutaraldehyde (Acros 233280250) in PBS for 1 hour. After washing with PBS, cells were permeabilized with 0.025% Triton X-100 (IBI Scientific, IB07100) in PBS for 10 minutes. Following additional PBS washes, cells were blocked in 3% BSA/PBS for at least an hour. Cells were treated with a 1:500 dilution of rat α-HA (Roche Sigma; 11867423001) and rabbit polyclonal antibodies against PyCITH or PyDDD for an hour and washed off with 3%BSA/PBS. Secondary antibodies of AlexaFluor 594-goat anti-rabbit (Thermo A11012) and AlexaFluor 633-goat anti-rat (Thermo A21094) were used at a 1:1000 dilution for 1 hour before washing in PBS. DAPI (Sigma D9564-10mg) was applied for nucleic acid staining at 1 µg/mL before fixing the cells in ProLong Glass (Invitrogen P36982) and sealing them on a slide with a coverslip and clear nail polish. Images were obtained on a Zeiss fluorescence microscope (Zeiss Axioscope A1 with an Axiocam 305 mono camera) using a 100x oil objective and processed in Fiji.

### RNA preparation and quantitative RT–PCR

Total RNA was extracted from parasites using TRIzol reagent (TransGen, cat#ET111-01-V2), and cDNA was synthesized using a reverse transcription kit (YoungGen, cat#24Y0007). For real-time quantitative PCR, the cDNA was diluted 20-fold with DEPC-treated water and subjected to SYBR Green quantification using Universal SYBR qPCR Master Mix (YoungGen, cat#QP101A) with gene-specific primers (Table S3). The 18S rRNA gene was used as an internal control. Three biological replicates, each with three technical replicates, were performed for each gene, and data were analyzed using the ΔΔCt method.

### RNA-seq and data analysis

Total RNA was extracted from purified *Plasmodium* gametocytes and then subjected to RNA-seq analysis (Suzhou, Genewiz). After assessing RNA integrity and purity by agarose gel electrophoresis and spectrophotometry, mRNA was enriched using Oligo(dT) beads (or by rRNA removal). cDNA was synthesized from the enriched mRNA using random primers, followed by end repair, poly(A) tailing, adapter ligation, PCR amplification, and bead purification. The constructed libraries were quantified and quality-controlled, then sequenced on an Illumina or MGI platform to generate 150 bp paired-end reads. Raw image data were converted to FASTQ format via base calling. Sequencing quality was evaluated with FastQC, and adaptor sequences, low-quality bases (quality value <20 or ambiguous ‘N’ bases) at the 5’ or 3’ ends, and reads shorter than 75 bp after trimming were removed using Cutadapt to obtain high-quality clean reads. The clean reads were aligned to the reference genome using Hisat2, and the percentages of total, uniquely mapped, and spliced reads were calculated; the distribution of reads across exonic, intronic, and intergenic regions was also examined. rMATS was used to identify and quantify five types of alternative splicing events (skipped exon, alternative 5’/3’ splice sites, mutually exclusive exon, and retained intron). Gene expression levels were calculated as FPKM values using HTSeq, and both read counts and FPKM values were exported for each gene. Differential expression analysis was performed using DESeq2, with |log₂ fold change| ≥ 1 and adjusted P ≤ 0.05 used as the thresholds for significance.

### Recombinant protein expression

A PCR product encoding ACDC-containing fragments (AP2-O3, residues 648-766; FD3, residues 1673-1773) and the LOTUS domain (GEP, residues 144-263) used as a negative-control bait were inserted into the pET-GST vector. The recombinant protein was expressed in *E. coli* BL21(DE3) (Coolaber, cat#CC553). Transformed bacteria were inoculated into LB medium containing 100 μg/ml carbenicillin (Sangon Biotech, cat#A600469) and cultured overnight at 37 °C with shaking at 220 rpm. The next day, the culture was diluted 1:50 into fresh LB medium and grown at 37 °C with shaking at 220 rpm until the OD600 reached 0.8–1.0. Protein expression was induced by adding isopropyl-β-D-thiogalactopyranoside (Coolaber, cat#CI6621) to a final concentration of 0.05 mM, followed by incubation at 16 °C with shaking at 140 rpm overnight. The recombinant protein was then affinity purified using glutathione resin (Biolinkedin, cat#PK-2006) according to the manufacturer’s instructions. The purification efficiency was assessed by SDS-PAGE and Coomassie Blue staining (Beyotime, cat# ST1119).

### GST pull-down and LC-MS/MS Analysis

Expression vectors encoding GST–AP2-O3–ACDC, GST–FD3–ACDC, and GST– LOTUS were transformed into *E. coli* BL21(DE3) (Coolaber, cat#CC553). For LC– MS/MS pull-downs, GST–AP2-O3–ACDC or GST–FD3–ACDC was used as the experimental bait, and GST–LOTUS was used as the negative-control bait; three biological replicates were analyzed per condition. Following induction, the bacterial cells were harvested, resuspended in an appropriate volume of bacterial lysis buffer, and ultrasonicated on ice. The lysate was centrifuged at 12,000 × g for 10 min at 4°C, and the supernatant was collected for use. Meanwhile, *6HA::ap2-o3; 4Myc::fd3* gametocyte proteins were extracted as prey. A 25 µl aliquot of glutathione resin (Biolinkedin, cat#PK-2006) was washed three times with pre-chilled wash buffer, mixed with the GST fusion protein supernatant, and incubated with rotation at 4°C for 3 h. After centrifugation, the supernatant was discarded, and the resin was washed three times with wash buffer. Subsequently, the *Plasmodium* gametocyte protein lysate was added to the resin-bound GST fusion protein and incubated with rotation at 4°C overnight. The resin was collected by centrifugation; the supernatant was discarded, and the resin was washed three times with pre-chilled wash buffer containing PMSF to remove non-specifically bound proteins. The resulting immunocomplexes were analyzed by SDS-PAGE and immunoblotting with corresponding antibodies, or by mass spectrometry (Shanghai Bioprofile Technology Company Ltd.) to identify proteins that potentially interact with the ACDC domain. Raw MS intensity values of 0 were imputed with half of the minimum intensity value. All intensities were subsequently log2-transformed, and fold changes were calculated between experimental groups.

### Co-immunoprecipitation

For co-immunoprecipitation (Co-IP), mouse blood samples from the *6HA::ap2-o3; 4Myc::fd3* line with high gametocytemia were harvested, and purified gametocytes were lysed in RIPA buffer containing protease inhibitors (RIPA:inhibitor = 100:1). After ultrasonication on ice (5 s on, 10 s off, 8 cycles) and incubation on ice for 30 min, the lysates were centrifuged at 12,000×g for 10 min at 4°C, and the supernatants were collected. A 25 µl aliquot of pre-equilibrated anti-Myc nanobody agarose beads (AlpalifeBio, cat#KTSM1364) was washed three times with pre-chilled dilution buffer, added to the supernatants, and incubated with rotation at 4°C for 3 h. After centrifugation, the beads were washed three times with dilution buffer. Finally, SDS-PAGE loading buffer was added to the beads; the mixture was heated at 100°C for 10 min, and the eluted samples were collected for Western blot analysis.

### Structure prediction and ligand library preparation

The three-dimensional structures of the *P. falciparum* AP2-O3 and FD3 ACDC domains were predicted using Protenix. The resulting models were inspected and prepared using PyMOL by removing water molecules and adding hydrogen atoms, followed by energy minimization using Rosetta Relax. For ligand library preparation, 276 compounds from the Tres Cantos Antimalarial Compound Set (TCAMS) and seven previously reported ACDC-targeting compounds were included, yielding a total of 283 compounds. Ligand structures were obtained in SMILES format, converted to three-dimensional conformers using RDKit with the MM2 molecular force field, and exported as PDB files.

### Identification of ligand-binding cavities and virtual screening

Putative ligand-binding cavities were identified using CB-DOCK2, and the highest-ranked cavity for each ACDC domain was selected for subsequent virtual screening. The PfAP2-O3 ACDC docking box was centered at x = −4.513, y = 4.222, and z = −5.692, with dimensions of 24 × 30 × 32 Å, whereas the PfFD3 ACDC docking box was centered at x = −0.132, y = −6.802, and z = −4.155, with dimensions of 26 × 24 × 24 Å. Docking was performed using AutoDock Vina 1.2.3. Ligands were converted to PDBQT format using Raccoon, and up to 20 binding modes were retained for each ligand. Compounds were ranked according to their predicted AutoDock Vina docking scores, and the top 20 compounds for each ACDC domain were retained for subsequent analysis.

### Protein–ligand interaction analysis

Predicted protein–ligand interactions of the three highest-ranked compounds for each ACDC domain were analyzed using the local version of PLIP. Interaction types, including hydrophobic contacts, hydrogen bonds, π-stacking, and halogen bonds, were identified from the predicted docking poses and visualized using PyMOL.

### Molecular dynamics simulations

The highest-ranked PfAP2-O3 ACDC–TCMDC-124223 and PfFD3 ACDC– TCMDC-134543 complexes were subjected to 350-ns all-atom molecular dynamics simulations using GROMACS 2023.3 [42]. Each system was solvated in a cubic TIP3P water box with a 10-Å boundary distance and neutralized with Na⁺ and Cl⁻ ions [43]. After energy minimization using the steepest-descent algorithm, each system underwent 100 ps of NVT equilibration followed by 100 ps of NPT equilibration at 310.15 K and 1 bar. During the simulations, hydrogen-containing bonds were constrained using the LINCS algorithm, electrostatic interactions were treated using the particle-mesh Ewald method, and a 1.2-nm non-bonded interaction cutoff was applied. Trajectories were corrected for periodic boundary conditions before analysis. Corresponding ligand-free ACDC domain simulations were analyzed using the same protocol for comparison.

### Trajectory analysis

Protein and ligand RMSD, residue-level RMSF, radius of gyration, protein–ligand hydrogen-bond occupancy, and trajectory-based protein–ligand interaction profiles were calculated using the processed molecular dynamics trajectories. RMSD analysis was used to assess ligand positional/conformational changes relative to the protein, whereas RMSF and radius of gyration were used to evaluate residue-level flexibility and overall structural compactness, respectively.

### MM/GBSA analysis

MM/GBSA calculations were performed on selected trajectory frames to estimate protein–ligand binding free-energy contributions [44–46]. The energy was decomposed into van der Waals, electrostatic, polar solvation, and nonpolar solvation components. Polar solvation energies were calculated using a generalized Born model with igb = 2 [47], whereas the nonpolar solvation contribution was estimated from solvent-accessible surface area using ΔGSA = 0.0072 × ΔSASA [48]. Entropic contributions were not included. Frames from 340–350 ns were analyzed for the PfAP2-O3 ACDC–TCMDC-124223 complex, whereas frames from 90–350 ns were analyzed for the PfFD3 ACDC–TCMDC-134543 complex.

### Antibodies and antiserum used in *P. yoelii*

The primary antibodies used were as follows: rabbit anti-HA (Cell Signaling Technology, cat# 3724; RRID: AB_1549585) for Western blot (1:1,000) and IFA (1:1,000); rabbit anti-HA (ABclonal, cat#AE036) for Western blot (1:5,000); mouse anti-Myc (Abmart, cat#M20002) for Western blot and IFA (1:1,000); mouse anti-GST (Abmart, cat#M20007) for Western blot (1:5,000); and mouse anti-α-Tubulin (Cell Signaling Technology, cat#3873S) for Western blot and IFA (1:1,000). The secondary antibodies used were: horseradish peroxidase-conjugated goat anti-rabbit IgG (Absin, cat#abs20040) and goat anti-mouse IgG (Absin, cat#abs20001), both at 1:5,000 dilution for Western blot; Alexa Fluor 555-conjugated goat anti-rabbit IgG (Thermo Fisher, cat#AB_141784), Alexa Fluor 488-conjugated goat anti-rabbit IgG (Thermo Fisher, cat#AB_10374301), Alexa Fluor 555-conjugated goat anti-mouse IgG (Thermo Fisher, cat#AB_141822), and Alexa Fluor 488-conjugated goat anti-mouse IgG (Thermo Fisher, cat#AB_2534069), all at 1:1,000 dilution for immunofluorescence. Hoechst 33342 (Thermo Fisher, cat#62249) was used for nuclear staining.

### Antibodies used in *P. falciparum*

The primary antibodies used were as follows: rat anti-HA (Roche Sigma; 11867423001); rabbit polyclonal antibodies against PyCITH and PyDDD [25, 26]. The secondary antibodies used were: Alexa Fluor 594-goat anti-rabbit (Thermo A11012) and Alexa Fluor 633-goat anti-rat (Thermo A21094). DAPI (Sigma D9564-10mg) was used for nuclear staining.

### Bioinformatics analysis and tools

Genomic sequences of target genes were downloaded from the PlasmoDB database (http://plasmodb.org/plasmo/). Single guide RNAs (sgRNAs) for each target gene were designed using the online tool EuPaGDT (http://grna.ctegd.uga.edu/). Sequence alignment was performed at https://espript.ibcp.fr/ESPript/ESPript/. PCR primers were designed using OligoArchitect™ Primer and Probe Design. The three-dimensional structures of the *P. falciparum* AP2-O3 and FD3 ACDC domains were predicted using the Protenix server (https://protenix-server.com/add-prediction). Putative active cavities of target proteins were identified using CB-Dock2. Statistical analysis was performed using GraphPad Prism (GraphPad Software).

## Data availability

The raw RNA-seq reads have been deposited in the NCBI SRA under BioProject accession PRJNA1494864 (https://www.ncbi.nlm.nih.gov/bioproject/?term=PRJNA1494864), and the processed gene expression matrix has been deposited in NCBI Gene Expression Omnibus under accession number GSE341438. The mass spectrometry proteomics data have been deposited in the ProteomeXchange Consortium via the PRIDE [49] partner repository with the dataset identifier PXD082017. All additional supporting data are included in the manuscript.

## Acknowledgements

We sincerely thank Denise Okafor for her thorough review of the manuscript and her insightful contributions to the modeling work. We also appreciate Jing Yuan and Bo Xiao for their careful review and valuable feedback. Additionally, we are grateful for the technical support from Phadcalc (www.phadcalc.com).

## Authors’ Contributions

**Jingjing Tang:** Conceptualization (with Z.L.), Investigation (generation of transgenic parasite strains, phenotypic analysis, RNA-seq profiling, GST pull-down assays, and protein-related experiments with Y.X.), Writing – review & editing. **Bing Liu:** Investigation (LC-MS/MS analysis), Formal analysis (bioinformatics analysis), Visualization, Writing – review & editing. **Aniko Verbrugge:** Investigation (experiments in *Plasmodium falciparum* with I.O.P.), Writing – review & editing. **Yangxin Xie:** Investigation (protein-related experiments with J.T.), Writing – review & editing. **Isadora Prata:** Investigation (experiments in *P. falciparum* with A.V.), Writing – review & editing. **Manuel Llinás:** Supervision, Project administration, Writing – original draft, Writing – review & editing. **Zhenkui Li:** Conceptualization, Supervision, Funding acquisition, Project administration, Writing – original draft, Writing – review & editing. All authors reviewed and approved the final manuscript.

## Competing interests

The authors declare no competing interests.

Declaration of generative AI and AI-assisted technologies in the writing process

During the preparation of this work, the authors used Grammarly (v1.2.291.1949) solely to polish the language and enhance the text’s readability and linguistic quality. After using this tool, the authors reviewed and edited the content as needed and take full responsibility for the publication’s content.

---

**Figure S1.**
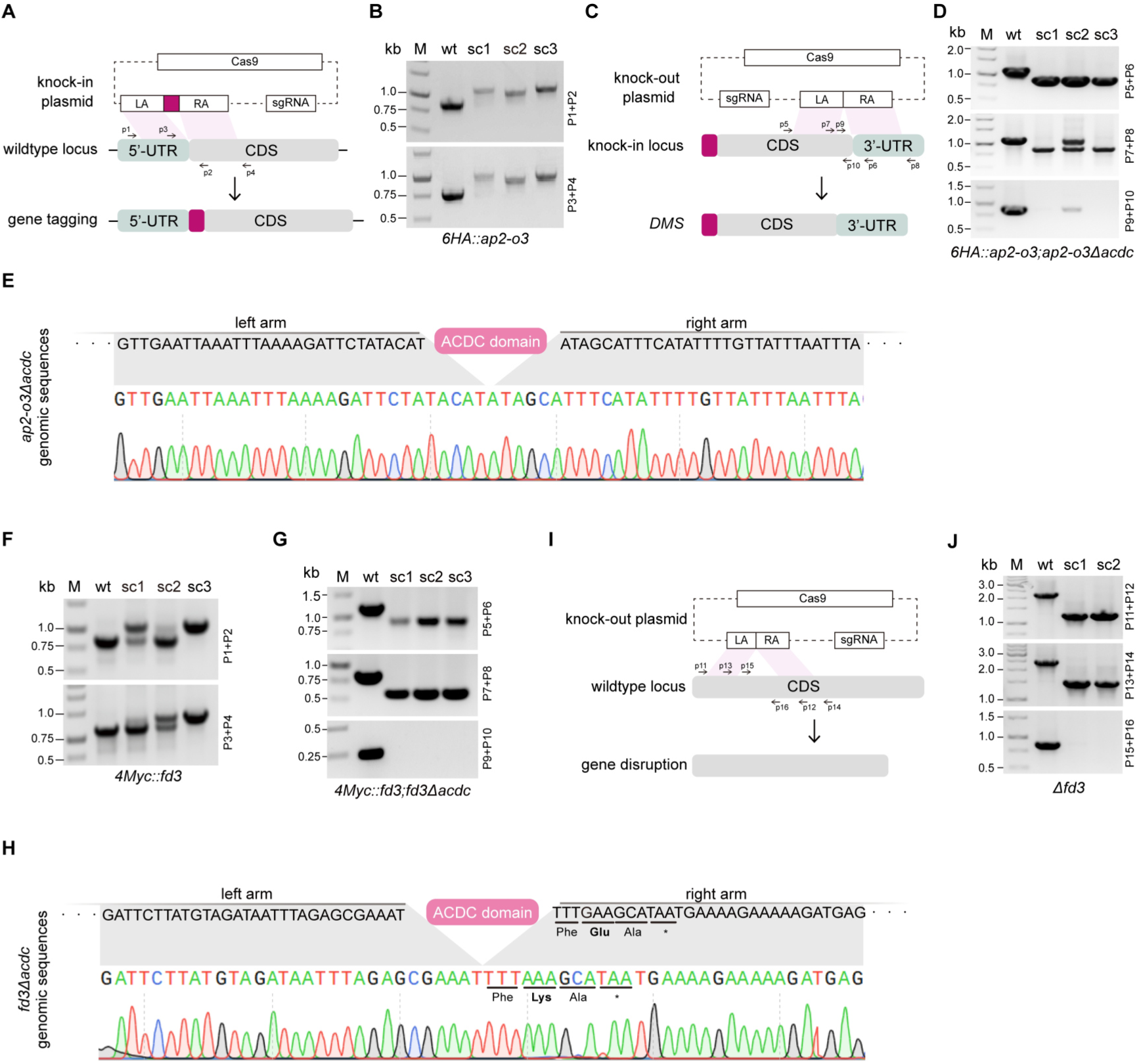
Generation and molecular validation of AP2-O3- and FD3-modified *P. yoelii* parasite lines. **(A)** Schematic showing how the endogenous target gene is tagged. **(B)** Diagnostic PCR verifies the insertion of the 6HA tag at the native *ap2-o3* locus. **(C)** Diagram illustrating the scarless, in-frame deletion of the ACDC-coding region. **(D)** Diagnostic PCR confirms the expected genomic arrangement of the *ap2-o3Δacdc* allele. **(E)** Sanger sequencing verifies the exact deletion of the AP2-O3 ACDC-coding sequence. **(F)** PCR validates the N-terminal 4Myc-tagged *fd3* parental line. **(G)** PCR confirms the *fd3Δacdc* allele. **(H)** Sanger sequencing verifies precise deletion of the FD3 ACDC-coding sequence. **(I)** Schematic approach for disrupting the coding region of *fd3*. **(J)** PCR validation of the *Δfd3* knockout line.

**Figure S2.**
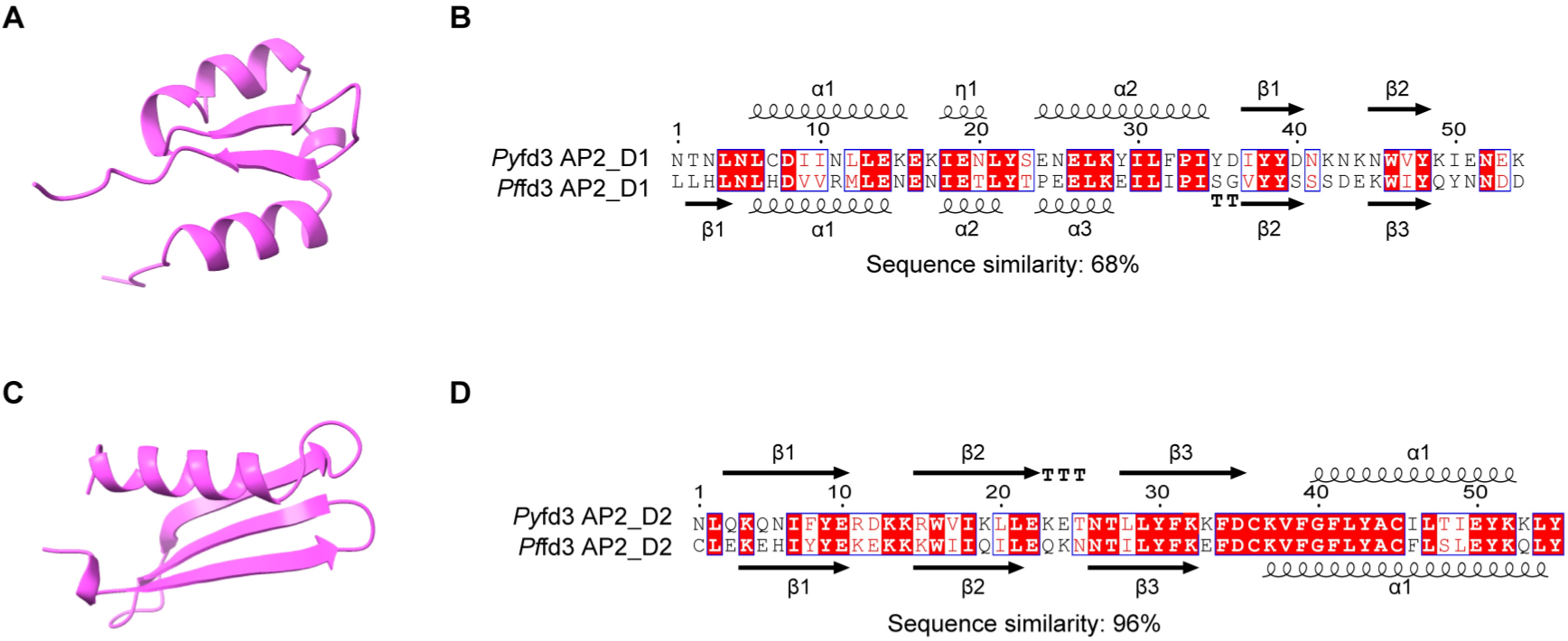
Sequence conservation and predicted structural features of the two candidate AP2-like domains in FD3. **(A, C)** Predicted three-dimensional structures of the AP2_D1 (residues 875–929) and AP2_D2 (residues 1202–1257) regions, respectively. AP2_D2 displays the characteristic fold of ApiAP2 DNA-binding domains, consisting of three antiparallel β-strands followed by an α-helix, this structural arrangement is less apparent in AP2_D1. **(B, D)** Cross-species sequence alignments of the corresponding AP2_D1 and AP2_D2 regions from *P. yoelii* (PyFD3) and *P. falciparum* (PfFD3), showing 68% and 96% sequence similarity, respectively. Predicted secondary-structure elements are indicated above and below the alignments, with α-helices represented by coils and β-strands by arrows. Identical and conserved residues are highlighted.

**Figure S3.**
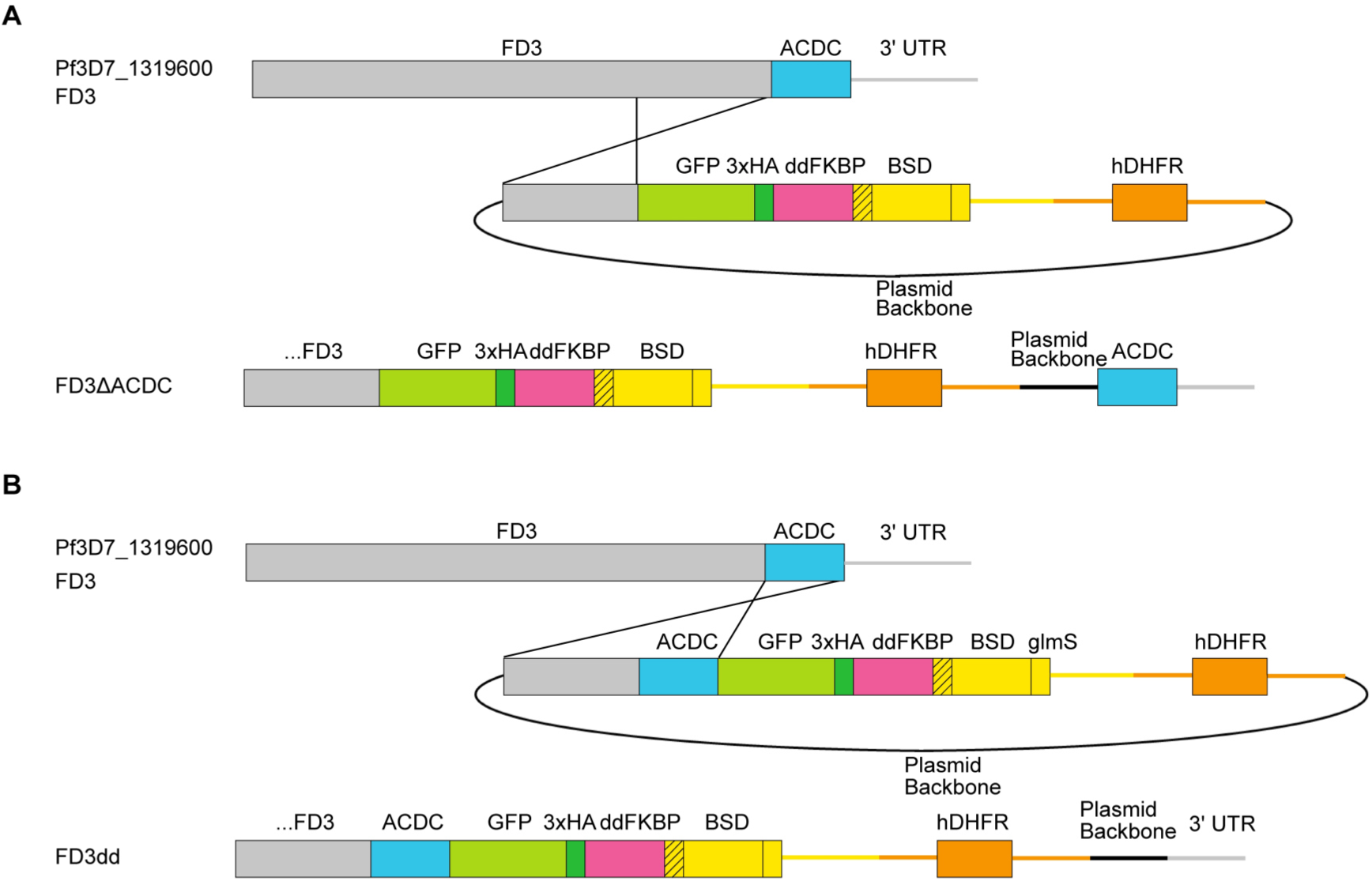
Construction of FD3-tagged and ΔACDC-tagged lines through single homologous recombination. The homology regions for FD3 were PCR-amplified and inserted upstream via restriction-enzyme cloning. These plasmids were transfected into the *P. falciparum* E5 strain.

**Figure S4.**
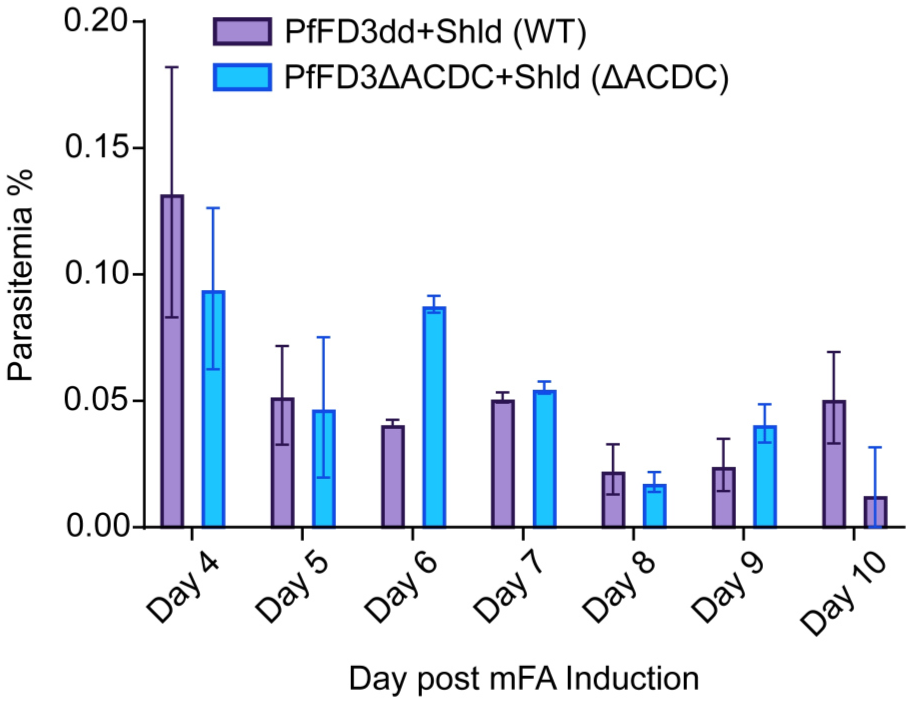
Gametocyte development of PfFD3dd and PfFD3ΔACDC parasites. Counts are normalized to parasitemia at Day 2 post-induction to account for growth differences before induction. N=2; At least 3000 RBCs were counted per day, per condition.

**Figure S5.**
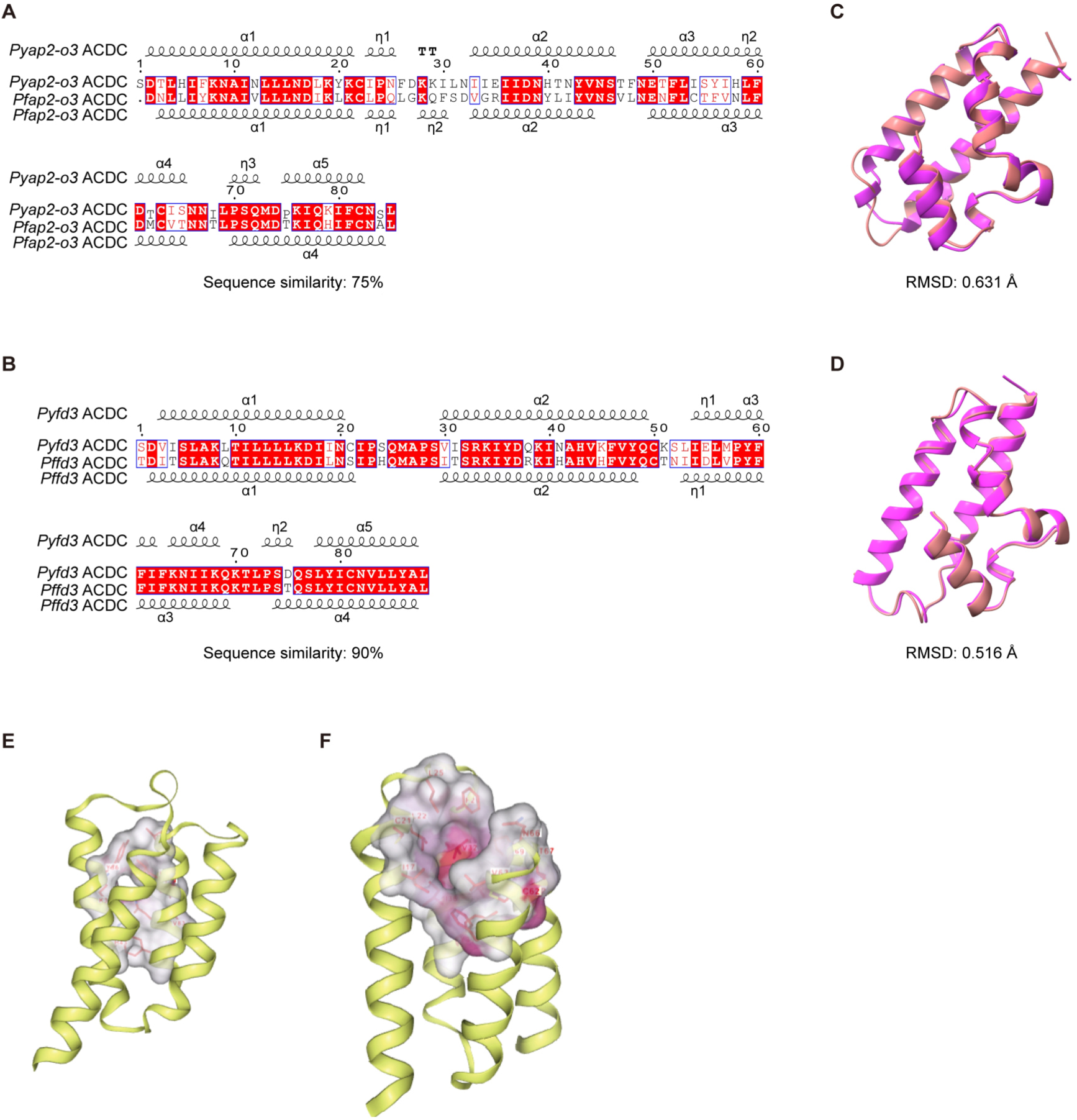
Sequence conservation, predicted structural conservation, and putative ligand-binding cavities of the AP2-O3 and FD3 ACDC domains. **(A)** Multiple sequence alignment of the AP2-O3 ACDC domains from *P. yoelii* (PyAP2-O3) and *P. falciparum* (PfAP2-O3). Identical residues are highlighted in red, and similar residues are boxed. Predicted secondary-structure elements are indicated above the alignment. **(B)** Multiple sequence alignment of the FD3 ACDC domains from PyFD3 and PfFD3. **(C)** Structural superposition of the Protenix-predicted PfAP2-O3-ACDC structure and the corresponding PyAP2-O3 structure. The backbone RMSD is 0.631 Å after removal of outlier atom pairs. **(D)** Structural superposition of the Protenix-predicted PfFD3-ACDC structure and the corresponding PyFD3 structure. The backbone RMSD is 0.516 Å after removal of outlier atom pairs. **(E)** Predicted ligand-binding cavity of the PfAP2-O3 ACDC domain identified using CB-DOCK2. **(F)** Predicted ligand-binding cavity of the PfFD3 ACDC domain identified using CB-DOCK2.

**Figure S6.**
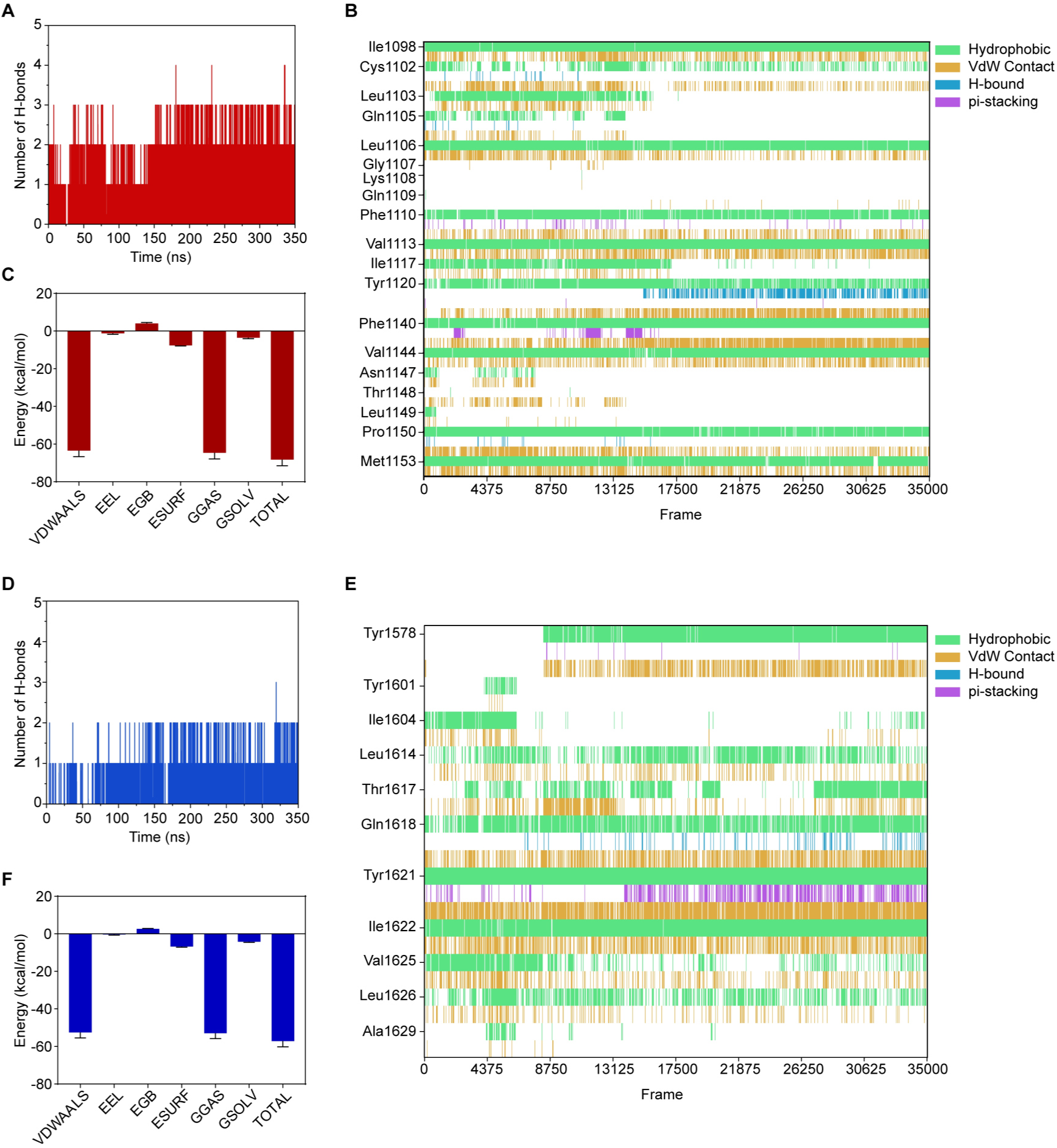
Hydrogen-bonding, residue-level interaction, and MM/GBSA analyses of the prioritized ACDC–ligand complexes during molecular dynamics simulations. **(A, D)** Number of protein–ligand hydrogen bonds as a function of simulation time for the PfAP2-O3 ACDC–TCMDC-124223 and PfFD3 ACDC–TCMDC-134543 complexes, respectively. **(B, E)** Frame-by-frame protein–ligand interaction profiles showing the persistence of hydrophobic, van der Waals, hydrogen-bond, and π-stacking interactions involving individual residues. **(C, F)** MM/GBSA energy decomposition for the indicated complexes. For the PfAP2-O3 ACDC–TCMDC-124223 complex, MM/GBSA analysis was performed using frames from 340–350 ns; for the PfFD3 ACDC–TCMDC-134543 complex, frames from 90–350 ns were analyzed. Interaction types are color-coded as indicated in the figure.

